# T7 DNA polymerase treatment improves quantitative sequencing of both double-stranded and single-stranded DNA viruses

**DOI:** 10.1101/2022.12.12.520144

**Authors:** Maud Billaud, Ilias Theodorou, Quentin Lamy-Besnier, Shiraz A. Shah, François Lecointe, Luisa De Sordi, Marianne De Paepe, Marie-Agnès Petit

## Abstract

**Background:** Bulk microbiome, as well as virome-enriched shotgun sequencing only reveals the double-stranded DNA (dsDNA) content of a given sample, unless specific treatments are applied. However, genomes of viruses often consist of a circular single-stranded DNA (ssDNA) molecule. Pre-treatment and amplification of DNA using the multiple displacement amplification (MDA) method enables conversion of ssDNA to dsDNA, but this process can lead to over-representation of these circular ssDNA genomes. A more recent alternative permits to bypass the amplification step, as library adapters are ligated to sheared and denatured DNA, after an end-modification step (xGen kit). However, the sonication step might shear ssDNA more efficiently than dsDNA, therefore introducing another bias in virome sequencing. These limitations prompted us to explore an alternative method of DNA preparation for sequencing mixed ssDNA and dsDNA viromes.

**Results:** Using a synthetic mix of viral particles, we made use of the T7 DNA polymerase (T7pol) to convert viral circular ssDNA molecules to dsDNA, while preventing over-replication of such molecules, as is the case with the Phi29 DNA polymerase. Our findings indicate that using T7pol and a mix of degenerated primers to convert ssDNA to dsDNA prior library preparation is a good alternative to the currently used methods. It better represents the original synthetic mixtures compared to MDA or direct application of the xGen kit. Furthermore, when applied to two complex virome samples, the T7pol treatment improved both the richness and abundance in the *Microviridae* fraction.

**Conclusion:** We conclude that T7pol pretreatment is preferable to MDA for the shotgun sequencing of viromes, which is easy to implement and inexpensive.

## Background

In the last ten years, many methodological improvements have helped generate informative metagenomics datasets for the virus-enriched fraction (or virome) of any type of environmental or biological sample (Castro-Mejia et al., 2015; Deng et al., 2019; Hsieh et al., 2021; Shkoporov et al., 2018; Wang et al., 2022). A particular difficulty of virome sequencing lies in the fact that the genetic material of viruses is not exclusively made of double-stranded (ds) DNA, as is the case for bacteria and higher organisms. It is frequently composed of single-stranded (ss) DNA and ss- or dsRNA. To our knowledge, a single method allowing the preparation of all these substrates at once for deep sequencing is not yet available. Even the simpler challenge of disposing of a method for the sequencing of both ds and ssDNA viruses is not completely solved.

Initially, most viromes were sequenced after a multiple displacement amplification (MDA) step, which was a convenient way to increase the usually low amounts of viral DNA to quantities compatible with library preparation. It became rapidly obvious that MDA introduced a bias favoring small and circular ssDNA viral genomes (the smallest ssDNA genomes are approximately 2000 nt long, while larger inoviruses are approximately 7000 nt long) (Kim and Bae, 2011; Yilmaz et al., 2010). This is because the Phi29 DNA polymerase used for MDA has potent strand displacement activity (Blanco et al., 1989). Therefore, once these small ssDNA genomes are converted into dsDNA, the polymerase proceeds for further rounds, generating long concatemers. Tests with synthetic mixes of viral genomes unambiguously demonstrated this caveat of MDA for mixes of ss- and dsDNA genomes (Roux et al., 2016).

Two main solutions were proposed to circumvent this problem. First, the “short MDA (sMDA)” method consisted of reducing the incubation time of the MDA treatment from the normally recommended 90- 120 minutes to 30 minutes (Deng et al., 2019). The second solution consisted of replacing the amplification step with a library kit able to add adapters to ssDNA molecules. Here, after an initial shearing step, all DNA molecule extremities, both ss- and dsDNA, receive a short extension using an “adaptase”, the nature of which is kept secret by the manufacturer. Then, an additional linker is added, allowing PCR amplification and sample labeling. This methodology is used in the Accel-NGS kit from Swift Biosciences (recently acquired by Integrated DNA Technologies (IDT) and rebranded as the xGen kit, below xGen). It was initially developed for the sequencing of ancient DNA, and has started to be used in the virome community (Roux et al., 2016). For this second method, tests with synthetic mixes showed that the bias in favor of ssDNA viruses was significantly reduced (Roux et al., 2016).

Using the xGen kit also comes with a drawback: the DNA shearing step is performed before the conversion of ssDNA to dsDNA, which may result in discrepancies if ssDNA is more readily sheared than dsDNA. This leads to possible downstream biases in library preparations, which require a homogeneous size distribution of the DNA fragments.

This prompted us to search for an alternative method of DNA preparation for the sequencing of mixed ssDNA and dsDNA viromes. We present here this new method based on the use of T7 DNA polymerase (T7pol) and compare it to two other methods: sMDA amplification, or no pretreatment. Then, all libraries were prepared with the xGen kit, for comparison purposes. We first compared the performance of the three methods on a synthetic community of five phages, and then tested the new T7Pol pre-treatment method followed by xGen libraries on two human viromes, in comparison with samples that had not been pre-treated.

## Methods

### DNA, enzymes, oligonucleotides, kits and other materials

ssDNA of M13mp18 and PhiX174, as well as dsDNA of PhiX174, were purchased from New England Biolabs (NEB, references N4040S, N3023L and N3021S, respectively). T7pol was purchased from NEB (ref. M0274) and used following the manufacturer’s instructions. Turbo DNAse I was purchased from Thermo Fisher Scientific (reference AM2222), and benzonase from Millipore (purity > 90%, reference 70746). For MDA treatments, the Genomiphi2 kit was purchased from Fisher Scientific (reference 10628505). The “xGen ssDNA Low-Input DNA Lib Prep 16rxn” (previously, Accel NGS S2 kit) was purchased from IDT (reference 10009859). Random oligonucleotides of a 20-nt size were purchased from Eurofins Genomics. SPRI Select magnetic beads were purchased from Beckman Coulter (reference B23318). To filter out bacteria from lysates and samples, polyethersulfone (PES) syringe filters 0.8/0.2 µm were used (Pall corporation, reference 4187).

### Phage lysates preparations

Two *Escherichia coli* infecting phages with ssDNA genomes were selected for the study: PhiX174 and an M13 derivative. PhiX174 is a *Microviridae* that forms a tail-less icosaedric capsid enclosing its circular 5386 nt ssDNA genome. It was grown on *E. coli* strain C, which codes for its appropriate core LPS receptor (Feige and Stirm, 1976). M13-yfp (De Paepe et al., 2010) is a filamentous phage of the *Inoviridae* family harboring a 7916 nt long circular ssDNA genome. It was grown on *E. coli* strain MG1655 F’kanR, a strain coding for the F pilus, which is the M13 receptor. Three dsDNA Caudoviricetes were also selected for the study: SPP1, T4 and lambda. T4 and lambda are *Escherichia coli*-infecting phages and were grown on *E. coli* strain MG1655. Inside the virions, the T4 genome is a 169 kb-long, linear dsDNA molecule, while the lambda linear genome has a shorter, 48 kb-long linear dsDNA genome. Finally, the *Bacillus subtilis* infecting phage SPP1 was propagated on strain YP886, generating virions containing a linear, 46 kb-long genome.

Phage lysates of T4, SPP1, lambda and phiX174 were prepared as described previously (Sausset et al., 2022), and bacterial remains were removed by centrifugation (5000 g, 7 min, 4°C), followed by filtration through PES filters. A lysate of M13-yfp was obtained by propagating the infected strain in 50 mL of LB-broth Lennox (Formedium) with 50 µg/mL kanamycin up to OD_600_ 1, removing bacteria by centrifugation (5000 g, 7 min, 4°C), and filtering the supernatant with PES filter. Stocks were kept at 4°C in salt-magnesium (SM) buffer (200 mM NaCl, 50 mM Tris pH 7.5, 10 mM MgSO4).

### Pure phage DNA preparation

From lysates, phages were concentrated by overnight centrifugation (20 000 g, 16 h), resuspended in 0.5 mL SM buffer, and loaded on SM-iodixanol gradients as described (Sausset et al., 2022). The phage-containing fraction was dialyzed against SM buffer and stored at 4°C. Prior to phage DNA extraction, purified particles were treated for 1 h at 37°C with DNaseI (8 units) and RNaseA (20 units) in SM buffer complemented with 0.5 mM CaCl2 to eliminate any residual bacterial DNA or RNA. The reaction was stopped by adding EDTA (20 mM).

To extract DNA from the capsids, samples were then treated with proteinase K (2 mg/mL) and SDS (0.5%) for 20 min at 56°C. Phage DNA was then extracted by phenol‒chloroform-isoamyl alcohol (25:24:1) extraction followed by two chloroform-isoamyl alcohol (24:1) purification steps. At each step, the solvent and aqueous phases were separated by centrifugation 15 minutes at 12 000g, 4°C. DNA was then supplemented with 300 mM potassium acetate pH 4.8, and precipitated with two volumes of ethanol, centrifuged for 30 min at 12000 g and 4°C, and the pellet was resuspended in 10 mM Tris buffer pH 8. Phage dsDNA concentration was estimated using a Qubit device.

### Phage quantification and preparation of the synthetic phage mixes

To prepare mixes containing phages in known concentrations, phage genome quantities of T4, lambda, SPP1 and PhiX174 were determined by qPCR, in comparison to dilutions of pure DNA standards. For these standards, concentration of PhiX174 ssDNA (from NEB) was verified both with a spectrophotometer (Nanodrop) and with a fluorescent-based method (Qubit). For M13 and PhiX174 ssDNA, Qubit values were 16 to 27% lower than Nanodrop values, respectively. Nanodrop values were kept, as they were closer to the Supplier’s indication (supplementary table 1) . For dsDNA qPCR standards, highly purified phage stocks of T4, SPP1 and lambda were prepared and quantified as described in the “Pure phage DNA preparation” section. Next, each stock of pure DNA was diluted in 10 mM Tris pH 8 to obtain a concentration of 5x10^6^ phage genomes in 6 μL. The samples were further diluted four times by 3-fold dilution steps, to obtain the calibration range (1.8x10^5^ to 5x10^6^ genomes), and aliquots stored at -20°C until use.

Before quantifying the four phage lysates, they were treated with DNaseI using 1 μL of enzyme/ml phage sample (1 h at 37°C) to eliminate any residual free DNA, and then treated at 95°C for 5 minutes, for capsid denaturation. Primers indicated in supplementary table 2 were used at a concentration of 200 nM. Three dilutions of each heat-denatured phage stock were compared to the calibration curves of highly purified DNA of the corresponding phage, as described above. qPCR was performed in a total volume of 15 μL (6 µL purified DNA or phage particle dilution and 9 µL of amplification mix) in MicroAmp Fast Optical 96-well plates sealed with MicroAmp Optical Adhesive Film (both from Applied Biosystems, reference 4346906 and 4311971, respectively) using the Luna universal qPCR master mix (NEB, ref M3003). Amplifications were run in duplicate on a StepOnePlus real-time PCR system with the following cycling conditions: 95°C for 5 min, (95°C for 15 s, 58°C for 45 s, 72°C for 30 s) for 45 cycles, 72°C for 5 min, 95°C for 15 s, 60°C for 15 s, and 95°C for 15 s. The analysis of the melting curves confirmed the specificity of the primers. Data analysis was performed with StepOne software 2.3.

The M13-yfp lysate could not be quantified by qPCR due to coat resistance to heat (González-Cansino et al., 2019), so it was quantified by plaque formation units on the MG1655 F’kan strain. Plaque counts of M13-yfp were easy to determine, as it is an M13mp18 derivative making much clearer plaques than the parental M13. Once each phage stock had been quantified, synthetic mixes were prepared. Initial phage concentrations in each stock, as well as volumes added to compose each mix, are presented in supplementary table 3.

### Synthetic mixes DNA extractions

To extract the DNA from the two synthetic mixes, an initial volume of 500 µL was used and DNA was extracted by phenol-chloroform treatment, as described in the “pure phage DNA preparation” section., resuspended in 25 µL of 10 mM Tris (pH 8), and dispatched twice for the three treatments as follows: 1 µL for MDA pretreatment (according to the Genomiphi2 kit instructions), 5 µL for T7pol pretreatment, and 5 µL for the xGen library without pretreatment. Final DNA yields were in average 100 ng for untreated and T7pol treated samples, 25 ng for the Genomiphi amplified mix with low proportion of ssDNA, and 500 ng for the Genomiphi amplified mix with high proportion of ssDNA.

### T7pol pretreatment

Primers for T7pol priming should have a minimum length of 15 nt (Hernandez et al., 2016), and their optimal size is 21 nt (C.C. Richardson, personal communication). To test whether degenerated primers could be used with T7pol to convert any phage ssDNA to dsDNA, degenerated primers of 20 nt were tested on ssDNA from PhiX174 and M13mp18. For the annealing step, samples containing 250 ng of M13mp18 and 250 ng of PhiX174 ssDNA in a final volume of 25 µL of water were incubated with varying concentrations of oligonucleotides for 5 minutes at 65°C (primer concentrations are indicated at this first annealing step, samples are then diluted 2-fold for the replication step). The samples were then allowed to cool down slowly by disposing them on floaters over a beaker containing 25 mL of 65°C water and placed at room temperature until final temperature was below 46°C (usually 10 minutes). Replication was then initiated by adding 300 µM dNTPs, 1X T7pol buffer, 50 µg/mL BSA and 5 units of T7pol in each tube completed with water to a final volume of 50 µL, followed by incubation for 5 minutes at 37°C. Reactions were stopped by placing the samples at 75°C for 20 minutes. Prior gel loading, samples were adjusted to 0.2% SDS, 10 mM EDTA and 0.5 mg/mL proteinase K, and incubated at 50°C for 30 minutes, to remove proteins that might prevent proper migration of the DNA molecules. Products (15 µL) were separated on a 0.7% agarose gel containing 40 µg/mL ethidium bromide. For the T7pol pretreatment of phage DNA mixes, the same conditions were applied, except that degenerated primers were used at a final concentration of 10 µM, together with 5 µL of DNA mix complemented with water to a final volume of 25 µL. The next replication step was performed in a final volume of 50 µL with 5 units of T7pol.

### Effect of sonication and magnetic bead size selection on ss- and dsDNA

PhiX174 ssDNA and dsDNA were each diluted at 20 ng/µL into 100 µL volumes and treated for sonication using conditions applied for library preparation (see below). Half of each sample was then submitted to SPRIselect magnetic bead binding, using conditions removing dsDNA fragments below 200 bp, as applied for library preparation (see below), except that elution was done in the same 50 µL volume of Tris 10 mM pH 8. To visualize ssDNA fragment sizes, products were then separated on a 10 cm-long polyacrylamide denaturing gel (5% of acryl/bis 19:1, 8 M urea, 1 x TBE). For this, 10 µL of formamide stained with bromophenol blue was added to 10 µL of each sample, incubated at 90°C for 5 minutes, and then placed on ice. Samples (20 µL) were then loaded on a preheated gel (>1 hour migration, 450 volts) and separated for ∼20 minutes at 300 volts until bromophenol blue reached the gel border. The gel was stained for 30 minutes with Syber Gold (Invitrogen, reference S11494, 1X final concentration), and bands were revealed by a Bio-Rad ChemiDocTM MP imaging system. The size at the maximum signal intensity of each smear was estimated with Image Lab software (Bio-Rad Laboratories). Six PCR fragments 601, 440, 330, 279, 200 and 116 bp-long were denatured with formamide and run as a size ladder.

### Library preparations, synthetic phage mix sequencing and analysis

DNA preparations from the synthetic phage mixes were first pretreated (or not) for the conversion of ssDNA to dsDNA using either the Phi29 polymerase of the Genomiphi2 kit according to the supplier’s instructions with a 30-minute incubation or T7pol (see Results section). Next, all samples were sonicated (100 µL in 1.5 mL tubes) with a Diagenode Bioruptor Pico device in a cooled water bath (4°C) for 21 sec with 90 sec breaks for 3 cycles, centrifuged to collect droplets on the side of the tubes, and sonicated again for 21 sec with 90 sec breaks for 2 cycles to generate DNA segments of an average size of approximately 500 bp. DNA samples were then concentrated to 20 µL using SPRIselect magnetic beads from Beckman Coulter (ratio beads/DNA=1.2) to remove fragments below 200 bp. All samples were then processed for library preparation using the xGen kit, following the manufacturer’s instructions. These library steps included (1) adaptase (37°C, 15 min then 95°C, 2 min); (2) extension (98°C, 30 s; 63°C, 15 s; 68°C, 5 min); (3) magnetic bead cleaning; (4) linker ligation (25°C, 15 min); (5) magnetic bead cleaning; (6) DNA concentration estimation by Qubit (high sensitivity kit); and (7) indexing by PCR (98°C, 30 s followed by 4 to 14 cycles of 98°C, 10 s; 60°C, 30 s; 68°C, 60 s) depending on the initial DNA concentration.

Paired-end sequencing (2x150 nt) was performed on an Illumina ISeq100 system at a depth of 800 000 reads per sample. Read cleaning was then computed with fastp (v 0.23.1, (Chen et al., 2018)) using the following parameters: --trimfront2 15 --trimtail1 2 --trimtail2 2 –trim_poly_g –trim_poly_x –r –W 4 – M 20 –u 30 –e 20 –l 100 –p –w 12. These parameters were used to remove the adapters present at read extremities with xGen libraries. Read mapping of both paired and unpaired reads on the phage genomes was then conducted with bwa (Li and Durbin, 2009) and the –a parameter. Next, the data were treated and analyzed with msamtools and the following parameters: msamtools filter –b –u –l 80 –p 95 –z 80 –besthit –S and then msamtools profile –multi=proportional. The number of reads mapped per genome was normalized in relative abundance (rel) at the msamtools profile step using the default parameter. Rel are computed as follows: an “abundance” value of each genome is estimated by the number of reads mapped to that genome, divided by its length and normalized by the sum of abundances across all five genomes. The proportion of each phage within the sequenced sample was then estimated by this rel ratio.

To compare final compositions of the sequenced mixes, Aitchison distances were used. This distance is a Euclidean distance based on centered log ratio (CLR) transformations of a virus count matrix. CLR transformation accounts for the compositionality that is especially pronounced when only a few taxa are present. In such cases Aitchison distances provide the most accurate metric for gauging the similarity of samples.

### Human fecal virome sample preparation and treatment

Fecal samples (0.2 g) from two healthy donors S4 and S18 were first resuspended in 4 mL of 10 mM Tris, pH 7.5, and then centrifuged (5 200 g, 30 min, 4°C). The supernatant was filtered using 0.2 µm diameter PES filters. Then, a benzonase-nuclease treatment (75 U per sample) was performed for 2 h at 37°C. The viral samples were then concentrated by precipitation overnight at 4°C using polyethyleneglycol (PEG 8000, 10% w/v) and NaCl 0.5 M. After centrifugation (5 200 g, 1 h, 4°C), the pellet was retrieved and processed for DNA extraction as described in the “Pure phage DNA preparation” section. Each DNA sample was then divided into two parts: one was kept unprocessed, and the other was further treated with T7pol as described above. All samples libraries were then prepared with the xGen kit. Paired-end sequencing (2x150 nt) was performed on an Illumina platform at a depth of 5 million read pairs per sample.

### Bioinformatics analysis of the virome samples

The different steps of the analysis are summarized in supplementary figure 1. First, reads were cleaned with fastp as described for the synthetic mix sequencing, above. To remove contamination, all the reads mapping to the human genome (GRCh38) according to Bowtie2 (v 2.4.1, (Langmead and Salzberg, 2012)) with the --very-sensitive option were then excluded. The remaining reads were assembled using SPAdes (options --only assembler -k 21,33,55,77,99,127, v 3.15.3, (Prjibelski et al., 2020)). Deduplication of contigs at the species level (95% average nucleotide identity) was performed using BLAT (Kent, 2002) as previously published (Shah et al., 2023). Viral contigs were identified through two parallel approaches: using softwares and searching by alignment against Refseq, as well as recently released phage gut databases. For the software, VirSorter2 (options --include-groups dsDNAphage,NCLDV,RNA,ssDNA,lavidaviridae --exclude-lt2gene --min-length 2000 --viral-gene-required --hallmark-required --min-score 0.9 --high-confidence-only, v 2.2.3, (Guo et al., 2021), VIBRANT (default parameters, v 1.2.1, (Kieft et al., 2020)), and CheckV (v 0.8.1, database v 1.5, (Nayfach et al., 2021a)) were used. For CheckV, only the contigs with a minimum of “medium” quality were considered viral. Contigs classified as viral by any of these three softwares (with the mentioned parameters) were kept. For the databases, contigs were aligned using BLAT (v 36, (Kent, 2002)) against the following databases: COPSAC (Shah et al., 2023), GVD (Gregory et al., 2020), MGV (Nayfach et al., 2021b), GPD (Camarillo-Guerrero et al., 2021) and viral RefSeq. The contigs that aligned against any contig in the databases with an alignment length >= 80% of the database contig were considered viral. A list of unique viral OTUs was generated by combining the outputs of both of these approaches. Gene calling on viral contigs was performed with Prodigal (Hyatt et al., 2010). *Microviridae* were identified among all these viral contigs by searching for the presence of either the major capsid protein (MCP) or replication initiation protein (Rep). For this, we ran HHsearch (Soding, 2005) against specific PHROGs profiles (PHROG 514 for the MCP, and PHROGs 713 and 942 for Rep), and retained hits with at least 90% probability and 60% coverage of the target protein (Terzian et al., 2021). To retrieve a closer relative for the identified *Microviridae*, a vContact2 graph was constructed (Bin Jang et al., 2019), which included these contigs and INPHARED (March 2023 version) as reference database (Cook et al., 2021). This graph was then analyzed with graphanalyzer (Pandolfo et al., 2022). For comparison purposes, the four samples were then subsampled to 2 M paired-end reads per sample using seqtk (v 1.3, https://github.com/lh3/seqtk). The subsampled reads were then aligned against the set of viral contigs using BWA-mem2 (v 2.2.1, (Vasimuddin et al., 2019)) with the -a option. Then, using msamtools, reads were first filtered to only keep alignments > 80 bp with >95% identity over >80% read length and only keeping the best alignment for each read (options -l 80 -p 95 -z 80 –besthit). To exclusively work with contigs for which there is sufficient support of its presence in a given sample, only contigs for which the overall sequencing depth was >= 1 and at least 50% of the contig length was covered were kept (values computed using msamtools coverage –summary command). The relative abundance of each contig passing these criteria was determined by using the msamtools profile (options --multi=proportional --unit=rel). At the end of these treatments, sample S4 contained 269 viral OTUs when no T7pol treatment had been applied, and 181 viral OTU for the T7pol pretreatment sample. Sample S18 was richer, with 645 and 702 vOTU without and with T7pol treatment, respectively. Those relative abundances were finally analyzed using the R package phyloseq (v 1.38.0, (McMurdie and Holmes, 2013)).

## Results

### Comparing the fate of ssDNA and dsDNA during library preparations

A necessary step of library preparations involves the production of DNA fragments of a homogeneous size, either by enzymatic treatment or ultrasonic DNA shearing. Enzymatic treatment cannot target ssDNA, and ssDNA treatment by sonication may result in systematic lower sizes compared to dsDNA. Indeed, the xGen protocol mentions that fragmentation of ssDNA may result in shorter fragments and lower library yields. To quantify this difference, samples of dsDNA and ssDNA corresponding to the RF1 replicating form and the virion form of the phage PhiX174 genome, respectively, were sonicated in parallel (replicates performed on two different days) for the duration applied during library preparation and separated on a urea-denaturing 5% acrylamide gel. After sonication, a lower size of DNA strands was systematically observed for the ssDNA samples compared to the dsDNA samples (peak size at 316 versus 562 nt for replicate 1 and 235 versus 330 nt for replicate 2, Figure 1: ds and ss slots).

**Figure 1.**
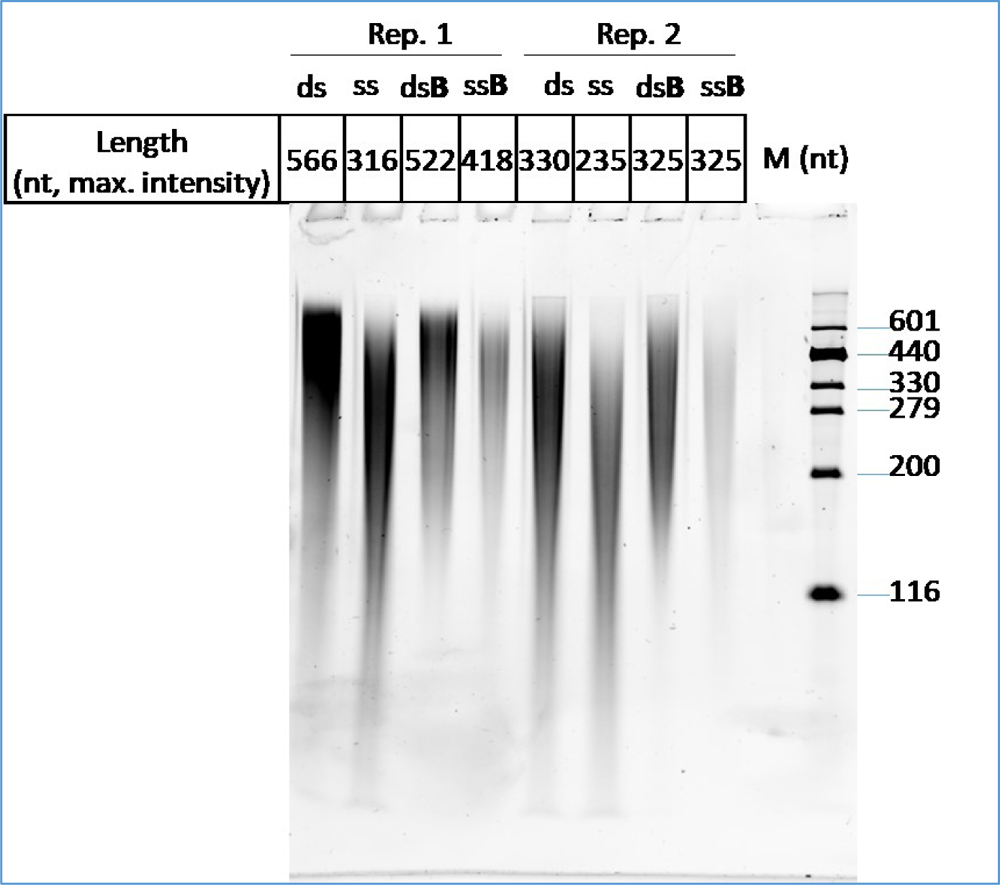
Effect of sonication and SPRI magnetic beads on ssDNA (ss slots) and dsDNA (ds slots) of PhiX174. All samples were sonicated for 5 cycles of 21 seconds, and a half volume was then treated with SPRI magnetic beads (B slots) to remove fragments below 200 bp. Samples were separated on a 5% acrylamide urea-denaturing gel. M: size marker. See the original gel picture in supplementary figure 2.

Some protocols concentrate the sonicated aliquots to 20 µL using magnetic beads for the adaptase step of the xGen kit (see Methods). We tested whether ssDNA fragments were preferentially lost at this step as well. Using the PhiX174 sonicated samples, total DNA recovery after the bead cleaning step was estimated with a Nanodrop apparatus. Whereas a total loss of 42-48% relative to input DNA was found for the dsDNA samples, this loss was more pronounced for ssDNA samples (74-84%). Migration of these samples next to their untreated controls on the denaturing gel indicated that this preferential loss was due to a higher size cutoff for ssDNA, compared to dsDNA, as the size at peak intensity increased after bead treatment for ssDNA (from 316 to 407 nt in replicate 1 and from 235 to 325 nt in replicate 2, Figure 1: ss and ssB labels), unlike for dsDNA (Figure 1: ds and dsB labels).

We concluded that both sonication and magnetic bead treatments had differential effects on ss- and dsDNA, with ssDNA being cut to 56-71% of the average size of dsDNA and small fragment removal by magnetic beads leading to a 2-fold lower recovery of ssDNA fragments compared to dsDNA.

#### Setting up a ssDNA-to-dsDNA conversion protocol using T7 DNA polymerase

To limit any bias in the treatments of ss- versus dsDNA molecules, we therefore searched for a quantitative method to convert ssDNA molecules to dsDNA prior to sonication. Contrary to the Phi29 DNA polymerase, T7pol has no intrinsic strand displacement activity, and does not open up the DNA strands ahead of polymerization process (Blanco et al., 1989; Canceill et al., 1999; Lechner and Richardson, 1983) (Figure 2A). Instead, a separate, replicative DNA helicase is responsible for this step, as in most living organisms (Canceill et al., 1999). As a consequence, once a first round of replication has been completed on a circular ssDNA molecule, T7pol stops and unloads, therefore preventing DNA overamplification.

**Figure 2:**
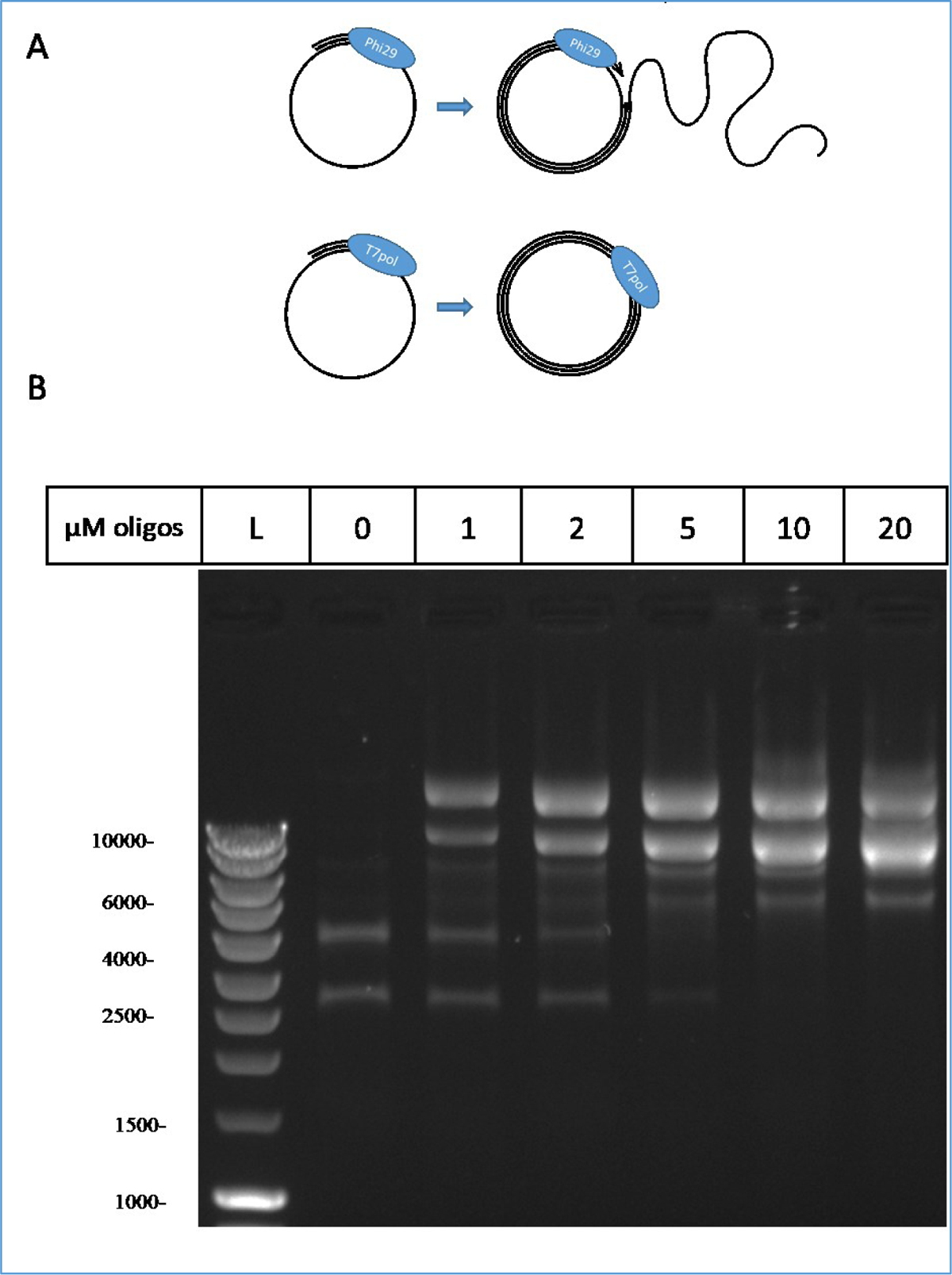
A. The different strand displacement activities of Phi29 DNA polymerase (up, rolling circle) and T7pol (down, stops after one round). B. Adjusting primer concentrations for T7pol conversion of ssDNA to dsDNA of M13 and PhiX174. Degenerated oligonucleotides (20 nt) were annealed at the indicated concentrations on a mix of M13 and PhiX174 ssDNA and incubated in the presence of T7pol. Products separated on 0.7% agarose gel are shown. Sizes (bp) of linear dsDNA standards (lane L, Smart ladder, Eurogentec) are indicated on the left (bp). See the original gel picture in supplementary figure 3.

To test conditions allowing T7pol replication initiation using degenerated oligonucleotides as primers, a 1:1 weight mix of PhiX174 and M13mp18 circular ssDNA molecules was used (10 µg/mL final DNA concentration). A range of oligonucleotide concentrations from 1 to 20 µM was tested, and repeated three times. A representative gel is shown Figure 2B. We found that 10 µM of 20 nt long degenerated oligonucleotides were sufficient to convert all the ssDNA species into dsDNA. In addition, and as expected, no smear indicative of further rounds of replication beyond the first one was detected. The marked intensity of the dsDNA bands, relative to ssDNA, was due to the higher affinity of ethidium bromide for dsDNA.

We concluded that T7pol treatment of circular and ssDNA should be an efficient and quantitative way to convert all DNA molecules to dsDNA prior to library preparation.

### A T7pol pretreatment of synthetic viromes generates more accurate results

We therefore proceeded to test this T7pol treatment on mock phage mixes and compared it to other ways of treating DNA for library preparation. For this, two synthetic mixes were prepared from stocks of five phages, two of which were ssDNA phages (M13-yfp and PhiX174), and three dsDNA phages (T4, lambda and SPP1). One mix contained 48.4% of ssDNA phage genomes (45.3% of M13-yfp, 3.1% of PhiX174), while the other contained only 5.9% of ssDNA phage genomes (5.5% M13-yfp and 0.4% PhiX174). The final and global phage concentration in each mix was approximately 10^9^ genomes/mL to mimic realistic viral concentrations in fecal samples (assuming a 10-fold dilution of fecal matter at 10^10^ particles/g).

Next, DNA was extracted from the two mixes, and each was subjected to one of the three following treatments (Figure 3A): (i) short MDA (sMDA), (ii) T7pol-dependent DNA replication using an excess of 10 µM degenerated oligonucleotides, and (iii) no pretreatment. Then, all samples were sonicated in a final volume of 100 µL and concentrated using magnetic beads. Finally, all were processed with the xGen kit for library preparation (see Methods for details). Each treatment was performed in duplicate, and the 12 samples were then sequenced on an Illumina platform (127 000 to 2.7 million reads per sample were obtained). Read mapping on the 5 phage genomes was converted into relative abundances, to estimate the proportion of each phage in the mix at the end of all treatments and to compare them to their theoretical initial ratio (Figure 3B, and supplementary table 4).

**Figure 3.**
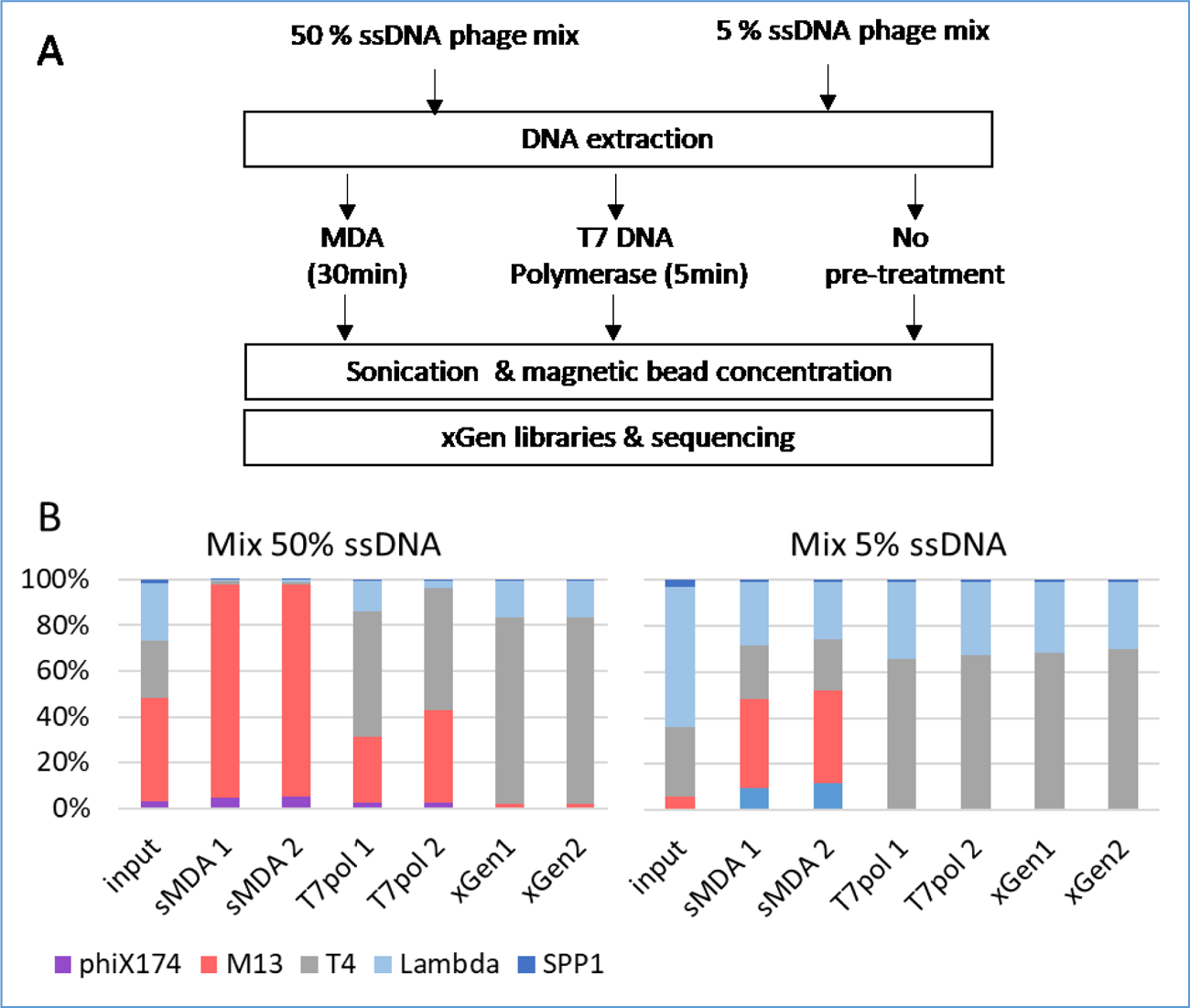
A: Comparison of protocols for sequencing ss- and dsDNA mixes. B. Input bar: Theoretical initial proportions of the 5 phages. Next six bars: Observed proportions of the 5 phages after deep sequencing (in duplicate, 1 and 2) of xGen DNA libraries either pre-treated with sMDA, T7pol, or without pre-treatment (xGen). Relative abundances are shown (rel values, see Methods).

We found that the DNA samples treated directly with xGen libraries (absence of pretreatment) led to a major depletion of the ssDNA phages: starting with the 48.4% input ssDNA mix, ssDNA phages only contributed 1.8-1.9% of the final sequencing result. The reduction of the ssDNA mix was even more drastic with 5.9% input ssDNA, as no reads matching the two ssDNA phages were obtained in one of the replicates, and M13 contributed only 0.09% of the sequenced virome in the other experiment. Since input DNA strands for library preparation were twice less abundant for ssDNA phages relative to dsDNA phages, the overall ssDNA phage depletion was 10-fold for the high input mix and over 30-fold for the low input mix.

With the short MDA pretreatment, starting with the 48.4% input ssDNA mix, 98% of ssDNA phage genomes were recovered, and 48-52% were recovered when using the 5.9% input ssDNA mix (here and below, no correction for single-strand molecules was applied, as all molecules were in the dsDNA state for library preparation). We conclude that overamplification occurred in both cases by factors of 2 and 8.6, respectively.

Finally, the T7pol pretreatment gave a result that was more congruent with input DNA: for the 48.4% ssDNA input mix, similar proportions (31-43%) of ssDNA phage were recovered, and 8.2-fold lower ssDNA (0.7%) was found for the 5.9% input mix.

To quantify the congruence between the initial mix compositions and the different pretreatments while accounting for the compositionality of the data, we computed Aitchison distances, a metric used to compare ecosystem compositions (Aitchison, 1982). Distances obtained between each input DNA mix and the various outputs, using the abundances from supplementary table 4, are reported in Table 1. T7pol pretreatment was more quantitative than both other methods, regardless of the input ssDNA/dsDNA ratio. Among the two other methods, short MDA pretreatment was more quantitative than direct xGen libraries for the mix with a low ssDNA concentration. However, the opposite was true at higher ssDNA concentrations. As a reference, the distance between the two input mixes themselves was also computed and found to be 3.05. This indicates that the T7pol pretreatment gave results closer to its initial mix (a distance of approximately 2) than the distance between the two input stocks.

**Table 1:** A. Aitchison distances between the compositions obtained from the different pretreatments and the input composition with low ssDNA amounts (5.9%). B. Same calculations for the high ssDNA input mix.

| A. | Mix low<br>ssDNA | B. | Mix high<br>ssDNA |
| --- | --- | --- | --- |
| xGen 1 | 8.17 | xGen 1 | 3.72 |
| xGen 2 | 5.97 | xGen 2 | 3.93 |
| sMDA 1 | 3.86 | sMDA 1 | 4.39 |
| sMDA2 | 4.02 | sMDA 2 | 4.34 |
| T7pol 1 | 2.29 | T7pol 1 | 2.15 |
| T7pol 2 | 2.23 | T7pol 2 | 2.31 |

### T7pol treatment of two viromes

To investigate the effect of T7pol treatment on complex environmental samples, two fecal samples were collected from healthy individuals (S4 and S18). Viromes were prepared, DNA extracted, and each sample was then treated or not with T7pol prior to sequencing, following the protocol described above. xGen libraries were then prepared exactly as for the synthetic communities. Reads were assembled de novo in contigs, which were then processed for the identification of viral ones (supplementary table 5). Among them, *Microviridae* were recognized by the presence of either the major capsid protein-encoding gene or the replication gene (see supplementary table 6 and supplementary figure 5 for details on these contigs). Mapping back the reads on all viral contigs allowed to determine the relative abundance of *Microviridae*. For the first S4 sample, the relative abundance of *Microviridae* increased from 24% to 74% of all viral contigs upon T7pol treatment. Richness in *Microviridae* also slightly increased, from 28 to 33 species. We noted, however, that 6 of the *Microviridae* contigs detected in the untreated samples were not present in the T7pol sample, while conversely, 11 *Microviridae* contigs were found only in the T7pol sample. A detailed analysis of the *Microviridae* profile indicated that each microvirus contig gained some level of abundance (some 8 +/- 6-fold on average), as indicated in Figure 4A (see also supplementary table 7). For the second S18 sample, the relative abundance of microviral species among viral contigs increased from 25% to 30%, and richness shifted from 5 to 8 species (Fig. 4B and supplementary table 7). We therefore conclude that the T7pol treatment improves overall the detection of *Microviridae* in complex samples, marginally in one case, and more substantially in the other case. Conversely, relative abundances of dsDNA, Caudoviricetes contigs were reduced upon T7pol treatment in sample S4. Their abundance was mostly unchanged in sample S18 (supplementary figure 4 and supplementary table 8). This was expected, as these two categories represent most of the signal, and therefore mirror each other in human fecal samples. We conclude that the T7pol treatment allows for better detection of ssDNA phages in natural samples, in agreement with the results obtained for synthetic communities.

**Figure 4:**
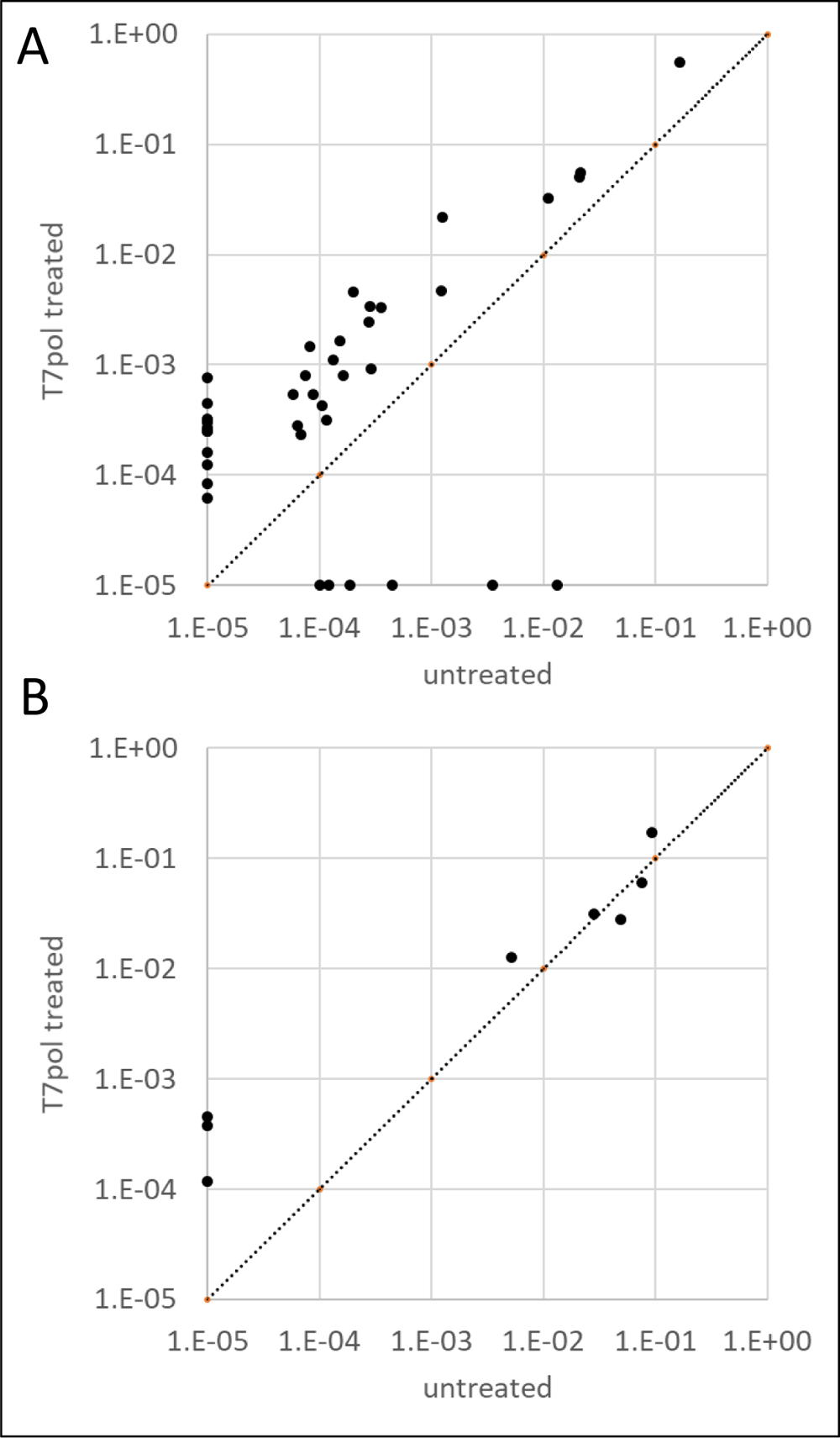
Relative abundance of each *Microviridae* detected in samples S4 (panel A) and S18 (panel B). Each dot represents one contig, with its abundance relative to all phages of the sample when sequenced without prior treatment for ssDNA x axis) or after T7pol treatment (Y axis). The dotted line shows the bisector. Contigs undetected with one of the treatments were assigned an artificial value of 10^-5^.

## Discussion

In the present benchmarking experiment, xGen libraries performed on untreated sonicated DNA were the most detrimental to ssDNA sequencing: ssDNA was recovered at levels 10 to over 30-fold less than expected. A previous study that also used synthetic mixes to compare MDA and xGen methods concluded that ssDNA sequencing was quantitative with xGen libraries (Roux et al., 2016). Many subtle differences might have led to the different conclusions. First, in this previous report, phage concentrations were estimated by epifluorescence, a method that might have underestimated the two *Microviridae* phage concentrations (PhiX174 and Alp3), as these particles are particularly dim under the microscope. Second, the kit used for DNA extraction (QIAamp DNA Mini Kit) was said to recover only 27% of the DNA for dsDNA phages while providing full recovery of ssDNA phages, so that proportions of dsDNA phages were artificially inflated *a posteriori*. In the present work, a phenol‒ chloroform extraction protocol was used, and no *a posteriori* corrections were made. Third, we found that ssDNA was more sensitive to shearing during the sonication step, and the parameters are likely to vary between sonication procedures. Fourth, we showed that some ssDNA was lost in our protocol at the magnetic bead step after sonication (the details of library preparation are not mentioned in (Roux et al., 2016). Clearly, all steps of DNA treatments should be carefully tested to ensure proper recovery of ssDNA. Nevertheless, our observation confirms the notion that pretreatments permitting the conversion of all ssDNA molecules to dsDNA prior to sonication should limit downstream bias, such as those we observed here. Having all DNA in the dsDNA state at the onset of library preparation also permits more accurate DNA concentration measurements, which is key for the tuning of sonication parameters, according to each kit supplier’s instructions.

Historically, MDA was the first method used to convert ssDNA to dsDNA; however, this method has long been known to be detrimental for the quantitative assessment of viromes containing a mix of ss- and dsDNA genomes (Kim and Bae, 2011; Roux et al., 2016). The standard 90-120 minute incubation time used for MDA was reported to amplify ssDNA by an excess of 33 to 44-fold when the input ssDNA was 2% (Roux et al., 2016). We found, however, that a shortening of the incubation time to 30 minutes (designated here “sMDA”) somewhat reduces the bias favoring ssDNA genomes: they were only in 8.6- fold excess when starting from 5.9% input ssDNA. However, starting with 48.4% ssDNA led to final ssDNA proportion of 98% of all sequenced DNA, somehow hiding all remaining dsDNA phage content.

T7pol pretreatment appears to be a better method than sMDA for sequencing DNA viromes. The overall distance between the final sequencing product and the input DNA was the least in this case, as well as being the most consistent for the two tested compositions. In particular, with respect to ssDNA, the final ssDNA genome proportions were only slightly lower than input ssDNA when the input phage mix had a high proportion of ssDNA (48.4%). However, for the low input ssDNA mix, the final proportion was 8.2-fold lower than expected. It may be that the annealing step required for T7pol- mediated DNA replication is less efficient when the ssDNA molecules are a minority within a complex sample. More work is needed to improve further the T7pol treatment. In fact, the optimal concentration of degenerated 20-mer oligonucleotides was very high (10µM), and even at this high level, only a minor fraction of these 20-mers are predicted to anneal perfectly with the template. We surmise that using shorter degenerated primers (such as 10-mer or 15-mer, instead of 20-mer) may function as well, and with lower overall primer concentrations. With the T7pol treatment, we also observed an improvement in the detection of *Microviridae*, which have ssDNA genomes, in two human fecal viromes sequenced with xGen libraries. This shows that such a treatment is feasible on natural samples and reinforces the interest in such a method.

It should be noted, however, that none of the protocols tested allowed the generation of an accurate image of the relative proportions of the five input phages of the two synthetic communities. This should be kept in mind when applying ecological metrics such as diversity and richness to viral metagenomics data. Compared to bacterial metagenomics, ecological analysis based on viral metagenomics is still in its infancy.

The T7pol pretreatment adds only a marginal cost (∼1 euro per sample) to the library preparations compared to a Genomiphi MDA pretreatment (20 euros per sample). Moreover, it should be possible to apply it to samples planned for any kind of downstream library preparation kit for low DNA amounts, such as the Nextera or TruSeq DNA nano kits from Illumina, making it independent of the 25% more expensive xGen kit, which was used here for comparison purposes. Further analyses are needed to explore this possibility.

Furthermore, the T7pol method described here might be a useful addition in the future, for projects aiming at simultaneously describing the bacterial and viral components of their microbial ecosystems without missing ssDNA viruses. In fact, several large databases of phage contigs have already been generated thanks to the data mining of shotgun metagenomics (Camarillo-Guerrero et al., 2021; Gregory et al., 2020; Nayfach et al., 2021b; Tisza et al., 2021). Such studies rely on the sorting of viral contigs among all other microbial contigs based on their characteristics and gene contents (Guo et al., 2021; Kieft et al., 2020). While this method is certainly efficient in detecting prophages, as well as phage DNA replicating inside bacteria and possibly encapsidated DNA, ssDNA phage genomes will probably escape analysis, as no pretreatment step ensuring their conversion into dsDNA is performed at present (Camarillo-Guerrero et al., 2021; Nayfach et al., 2021b; Tisza et al., 2021). We noted, however, that *Microviridae* were sometimes present in studies assembling viral sequences from such shotgun metagenomic samples which, in theory, should not have sequenced ssDNA (Camarillo-Guerrero et al., 2021; Nayfach et al., 2021b; Tisza et al., 2021). This might be explained if dsDNA replication intermediates were extracted together with bacterial DNA. We cannot exclude, however, that, depending on the biochemistry of each library preparation kit, the ssDNA fraction might be included unknowingly, and sequenced quantitatively. In any case, the present work shows that adding a T7 polymerase pretreatment can improve the sequencing of ssDNA viruses.

## Conclusion

We conclude that T7pol pretreatment is a better alternative to MDA for the shotgun sequencing of both the dsDNA and ssDNA fractions of viromes, which is easy to implement and inexpensive.

## Declarations

*Ethics approval and consent to participate.* Approval for human studies was obtained from the local ethics committee (Comité de Protection des Personnes Ile-de-France IV, IRB 00003835 Suivitheque study; registration number 2012/05NICB).

## Consent for publication

Not applicable

## Availability of data and material

The supplementary material (containing all supplementary figures and tables), as well as the raw reads generated for this study are available at https://doi.org/10.57745/C0BAPR.

## Authors’ contributions

MB, IT, QLB and FL conceived and realized the experiments, analyzed the data and contributed to the redaction of the manuscript. MAP supervised the research and wrote the manuscript. MDP and LDS supervised the research and contributed to the redaction of the manuscript. SAS contributed to the statistical analysis of the data and to the redaction of the manuscript.

## Acknowledgments

The Migale platform at INRAE, on which bioinformatics analyses were conducted, is warmly thanked for its constant support and help.

---

**Supplementary table 1:** ssDNA concentrations (ng/µL), tested by two different methods.

|  | Nanodrop |  | Qubit |  |
| --- | --- | --- | --- | --- |
|  | measure 1 | measure 2 | measure 1 | measure 2 |
| ssDNA PhiX174 NEB, 100 ng/μL | 101.6 | 95 | 76.2 | 67.6 |
| ssDNA M13mp18 NEB, 25 ng/μL | 18.8 | 18.5 | 15.7 | 15.4 |
| Standard ssDNA of the Qubit kit, 20 ng/μL | 23.8 | 23.1 |  |  |

**Supplementary table 2:**
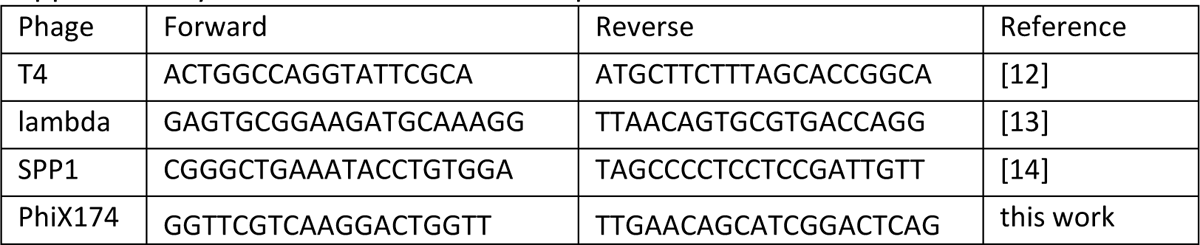
Primers used for qPCR.

**Supplementary table 3:**
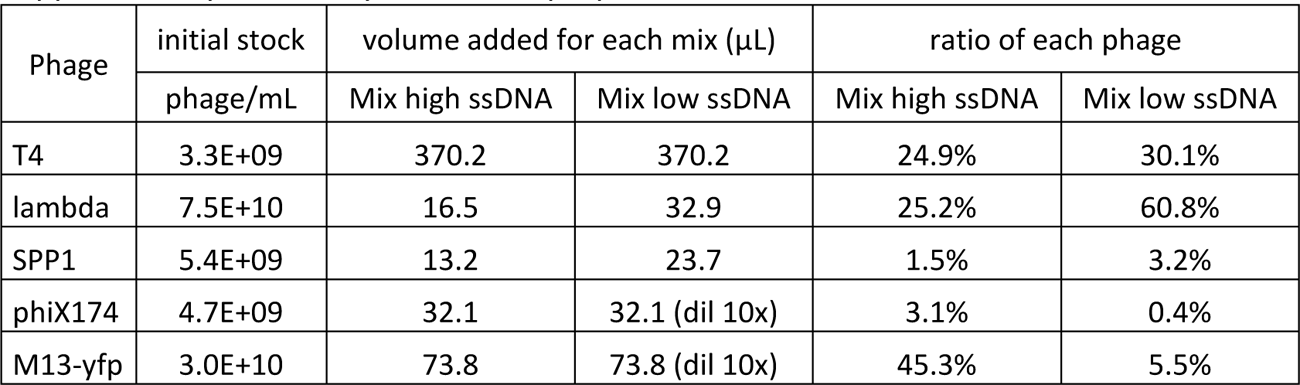
Synthetic mix preparations.

**Supplementary table 4:** Proportion of the five genomes in input mixes and after sequencing, using the different protocols. A panel, high ssDNA mix, B panel, low ssDNA mix.

| A. high ssDNA mix | input | short MDA 1 | short MDA 2 | T7pol 1 | T7pol 2 | xGen without pretreatment 1 | xGen without pretreatment 2 |
| --- | --- | --- | --- | --- | --- | --- | --- |
| T4 (ds) | 24.9% | 1.2% | 1.0% | 54.8% | 53.6% | 81.4% | 81.4% |
| Lambda (ds) | 25.2% | 0.8% | 0.9% | 13.4% | 2.9% | 16.2% | 16.2% |
| SPP1 (ds) | 1.5% | 0.0% | 0.0% | 0.5% | 0.7% | 0.6% | 0.6% |
| PhiX174 (ss) | 3.1% | 4.7% | 5.1% | 2.6% | 2.5% | 0.2% | 0.1% |
| M13-yfp (ss) | 45.3% | 93.3% | 92.9% | 28.6% | 40.2% | 1.7% | 1.7% |
| %ssDNA, in total | 48.4% | 98.0% | 98.0% | 31.3% | 42.7% | 1.9% | 1.8% |

| B. low ssDNA mix | input | short MDA 1 | short MDA 2 | T7pol 1 | T7pol 2 | xGen without pretreatment 1 | xGen without pretreatment 2 |
| --- | --- | --- | --- | --- | --- | --- | --- |
| T4 (ds) | 30.1% | 23.3% | 22.2% | 65.3% | 66.5% | 68.1% | 69.7% |
| Lambda (ds) | 60.8% | 27.5% | 24.9% | 33.1% | 31.9% | 31.0% | 29.5% |
| SPP1 (ds) | 3.2% | 1.0% | 1.0% | 1.0% | 1.0% | 0.9% | 0.7% |
| PhiX174 (ss) | 0.4% | 9.8% | 11.4% | 0.2% | 0.1% | 0.00% | 0.00% |
| M13-yfp (ss) | 5.5% | 38.4% | 40.3% | 0.5% | 0.6% | 0.00% | 0.09% |
| %ssDNA, in total | 5.9% | 48.2% | 51.7% | 0.7% | 0.7% | 0.0% | 0.1% |

**Supplementary table 5:** properties of the 4 assembled viromes.

|  | **Raw reads**<br>**(number of pairs)** | **Clean, non-human**<br>**reads (number of pairs)** | **Number of present**<br>**viral contigs** |
| --- | --- | --- | --- |
| **S4_Raw** | 3 944 094 | 3 101 396 | 645 |
| **S4_T7Pol** | 2 654 472 | 2 016 857 | 702 |
| **S18_Raw** | 47 423 047 | 36 124 946 | 269 |
| **S18_T7Pol** | 6 266 796 | 5 331 092 | 181 |

**Supplementary table 6:** properties of the microviruses found in samples S4 and S18.

| Contig | CheckV Quality | Length | GC% | Closer Relative | HostGenus |
| --- | --- | --- | --- | --- | --- |
| Microvirus_01 | High-quality | 5816 | 47.54 | Human gut gokushovirus | NA |
| Microvirus_02 | Medium-quality | 4309 | 38.8 | NA | Prevotella |
| Microvirus_03 | Medium-quality | 4289 | 38.7 | NA | Prevotella |
| Microvirus_04 | Complete | 5961 | 43.9 | Microviridae sp. | Faecalibacterium |
| Microvirus_05 | Complete | 5454 | 44.98 | Sigmofec virus UA08Rod_5824 | NA |
| Microvirus_06 | Complete | 7094 | 36.31 | Microvirus sp. | Prevotella |
| Microvirus_07 | Complete | 5880 | 43.59 | Poophage MBI-2016a | NA |
| Microvirus_08 | Medium-quality | 3185 | 36.17 | NA | NA |
| Microvirus_09 | High-quality | 6650 | 40.33 | Dipodfec virus UOA04_Rod_446 | NA |
| Microvirus_10 | High-quality | 5925 | 44.1 | Dipodfec virus UOA04_Rod_841 | NA |
| Microvirus_11 | Complete | 6229 | 40.78 | Microviridae sp. | Phocaeicola |
| Microvirus_12 | Complete | 6516 | 41.9 | Microviridae sp. | NA |
| Microvirus_13 | Complete | 5845 | 47.44 | NA | Alistipes |
| Microvirus_14 | Complete | 5063 | 43.33 | Gokushovirinae sp. | NA |
| Microvirus_15 | Medium-quality | 3198 | 36.59 | NA | NA |
| Microvirus_16 | Complete | 4891 | 42.32 | Capybara microvirus Cap1_SP_263 | NA |
| Microvirus_17 | Complete | 6300 | 38.59 | Microviridae sp. | Bacteroides |
| Microvirus_18 | Complete | 5873 | 45.79 | Gokushovirus WZ-2015a | Faecalibacterium |
| Microvirus_19 | Complete | 5847 | 43.66 | Gokushovirus WZ-2015a | NA |
| Microvirus_20 | Complete | 5758 | 43.4 | Gokushovirus WZ-2015a | Faecalibacterium |
| Microvirus_21 | Complete | 6720 | 37.35 | Dipodfec virus UOA04_Rod_499 | NA |
| Microvirus_22 | Medium-quality | 2631 | 43.18 | NA | NA |
| Microvirus_23 | Complete | 6037 | 37.22 | Microvirus sp. | NA |
| Microvirus_24 | Complete | 6123 | 35.72 | Microvirus sp. | NA |
| Microvirus_25 | High-quality | 6775 | 38.79 | Microvirus sp. | Prevotella |
| Microvirus_26 | Complete | 4800 | 45.02 | Human gut gokushovirus | NA |
| Microvirus_27 | Low-quality | 2125 | 41.88 | Gokushovirus WZ-2015a | NA |
| Microvirus_28 | High-quality | 5711 | 35.95 | Microvirus sp. | NA |
| Microvirus_29 | Medium-quality | 3568 | 47.76 | Human gut gokushovirus | NA |
| Microvirus_30 | Complete | 6963 | 38.33 | Human gut microviridae SH-CHD8 | NA |
| Microvirus_31 | Complete | 5168 | 44.54 | Microviridae sp. | NA |
| Microvirus_32 | Complete | 5176 | 41 | Gokushovirus WZ-2015a | NA |
| Microvirus_33 | Complete | 5499 | 46.79 | Dipodfec virus UOA04_Rod_999 | Alistipes_A |
| Microvirus_34 | High-quality | 5875 | 45.28 | Gokushovirus WZ-2015a | Faecalibacterium |
| Microvirus_35 | Complete | 5335 | 45.74 | NA | NA |
| Microvirus_36 | Complete | 5536 | 44.09 | Microviridae sp. | Dysosmobacter |
| Microvirus_37 | Medium-quality | 3381 | 38.15 | Tortoise microvirus 14 | NA |
| Microvirus_38 | Medium-quality | 3042 | 35.77 | Gokushovirinae sp. | NA |
| Microvirus_39 | Medium-quality | 2986 | 47.56 | Gokushovirus WZ-2015a | NA |
| Microvirus_40 | Complete | 5919 | 45.65 | Gokushovirus WZ-2015a | Faecalibacterium |
| Microvirus_41 | Complete | 5934 | 45.8 | NA | NA |
| Microvirus_42 | Complete | 6619 | 37.68 | Tortoise microvirus 1 | NA |
| Microvirus_43 | Complete | 5157 | 41.09 | Microviridae sp. | NA |
| Microvirus_44 | Complete | 5874 | 43.65 | Gokushovirus WZ-2015a | Faecalibacterium |

**Supplementary table 7:**
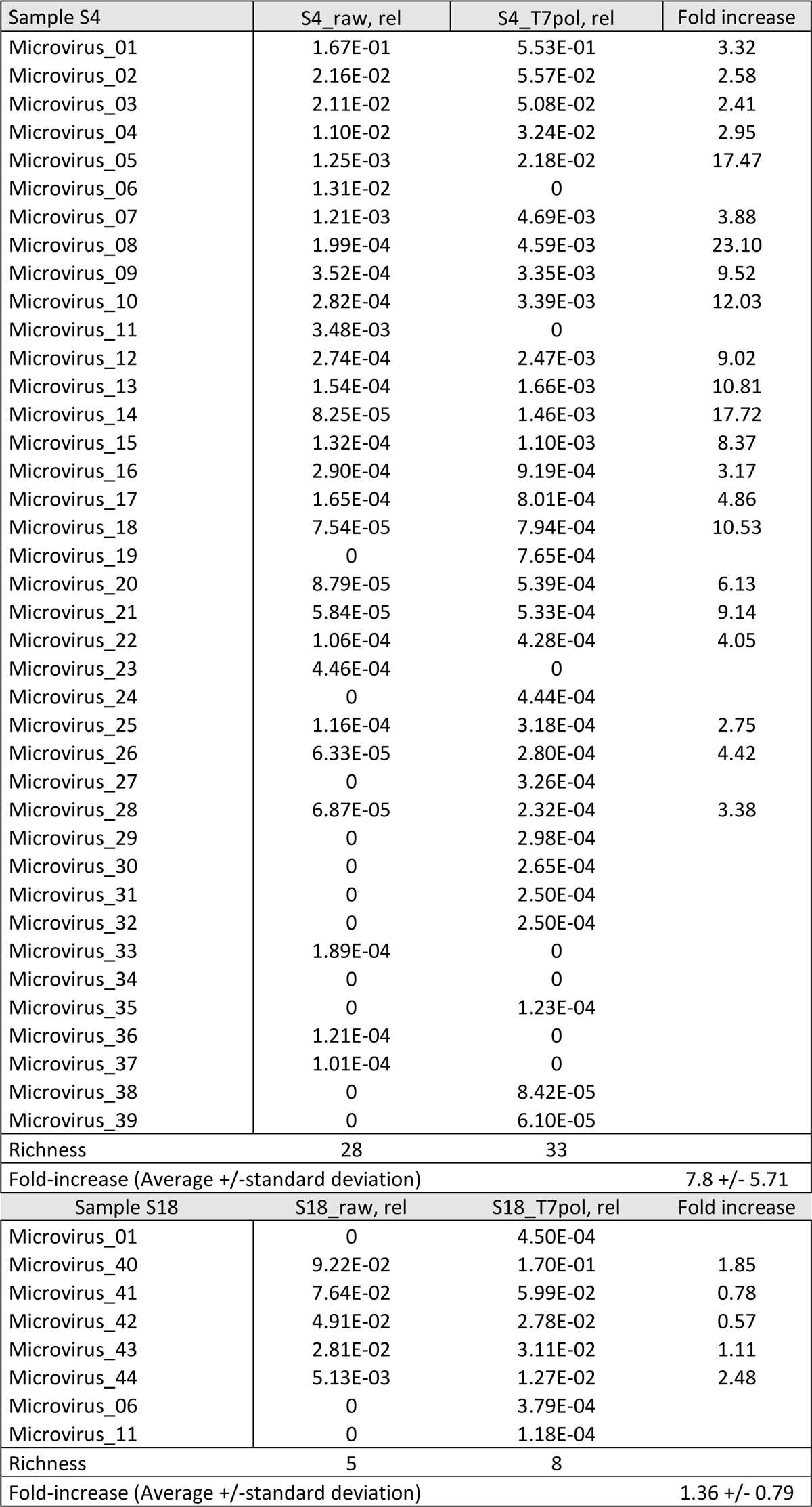
Relative abundance (rel) of each microviral contig in samples S4 and S18 before (“raw” column) and after T7pol treatment (“T7pol column”).

| Sample S4 | S4_raw, rel | S4_T7pol, rel | Fold increase |
| --- | --- | --- | --- |
| Microvirus_01 | 1.67E-01 | 5.53E-01 | 3.32 |
| Microvirus_02 | 2.16E-02 | 5.57E-02 | 2.58 |
| Microvirus_03 | 2.11E-02 | 5.08E-02 | 2.41 |
| Microvirus_04 | 1.10E-02 | 3.24E-02 | 2.95 |
| Microvirus_05 | 1.25E-03 | 2.18E-02 | 17.47 |
| Microvirus_06 | 1.31E-02 | 0 |  |
| Microvirus_07 | 1.21E-03 | 4.69E-03 | 3.88 |
| Microvirus_08 | 1.99E-04 | 4.59E-03 | 23.10 |
| Microvirus_09 | 3.52E-04 | 3.35E-03 | 9.52 |
| Microvirus_10 | 2.82E-04 | 3.39E-03 | 12.03 |
| Microvirus_11 | 3.48E-03 | 0 |  |
| Microvirus_12 | 2.74E-04 | 2.47E-03 | 9.02 |
| Microvirus_13 | 1.54E-04 | 1.66E-03 | 10.81 |
| Microvirus_14 | 8.25E-05 | 1.46E-03 | 17.72 |
| Microvirus_15 | 1.32E-04 | 1.10E-03 | 8.37 |
| Microvirus_16 | 2.90E-04 | 9.19E-04 | 3.17 |
| Microvirus_17 | 1.65E-04 | 8.01E-04 | 4.86 |
| Microvirus_18 | 7.54E-05 | 7.94E-04 | 10.53 |
| Microvirus_19 | 0 | 7.65E-04 |  |
| Microvirus_20 | 8.79E-05 | 5.39E-04 | 6.13 |
| Microvirus_21 | 5.84E-05 | 5.33E-04 | 9.14 |
| Microvirus_22 | 1.06E-04 | 4.28E-04 | 4.05 |
| Microvirus_23 | 4.46E-04 | 0 |  |
| Microvirus_24 | 0 | 4.44E-04 |  |
| Microvirus_25 | 1.16E-04 | 3.18E-04 | 2.75 |
| Microvirus_26 | 6.33E-05 | 2.80E-04 | 4.42 |
| Microvirus_27 | 0 | 3.26E-04 |  |
| Microvirus_28 | 6.87E-05 | 2.32E-04 | 3.38 |
| Microvirus_29 | 0 | 2.98E-04 |  |
| Microvirus_30 | 0 | 2.65E-04 |  |
| Microvirus_31 | 0 | 2.50E-04 |  |
| Microvirus_32 | 0 | 2.50E-04 |  |
| Microvirus_33 | 1.89E-04 | 0 |  |
| Microvirus_34 | 0 | 0 |  |
| Microvirus_35 | 0 | 1.23E-04 |  |
| Microvirus_36 | 1.21E-04 | 0 |  |
| Microvirus_37 | 1.01E-04 | 0 |  |
| Microvirus_38 | 0 | 8.42E-05 |  |
| Microvirus_39 | 0 | 6.10E-05 |  |
| Richness | 28 | 33 |  |
| Fold-increase (Average +/-standard deviation) |  |  | 7.8 +/- 5.71 |
| Sample S18 | S18_raw, rel | S18_T7pol, rel | Fold increase |
| Microvirus_01 | 0 | 4.50E-04 |  |
| Microvirus_40 | 9.22E-02 | 1.70E-01 | 1.85 |
| Microvirus_41 | 7.64E-02 | 5.99E-02 | 0.78 |
| Microvirus_42 | 4.91E-02 | 2.78E-02 | 0.57 |
| Microvirus_43 | 2.81E-02 | 3.11E-02 | 1.11 |
| Microvirus_44 | 5.13E-03 | 1.27E-02 | 2.48 |
| Microvirus_06 | 0 | 3.79E-04 |  |
| Microvirus_11 | 0 | 1.18E-04 |  |
| Richness | 5 | 8 |  |
| Fold-increase (Average +/-standard deviation) |  |  | 1.36 +/- 0.79 |

**Supplementary table 8:**
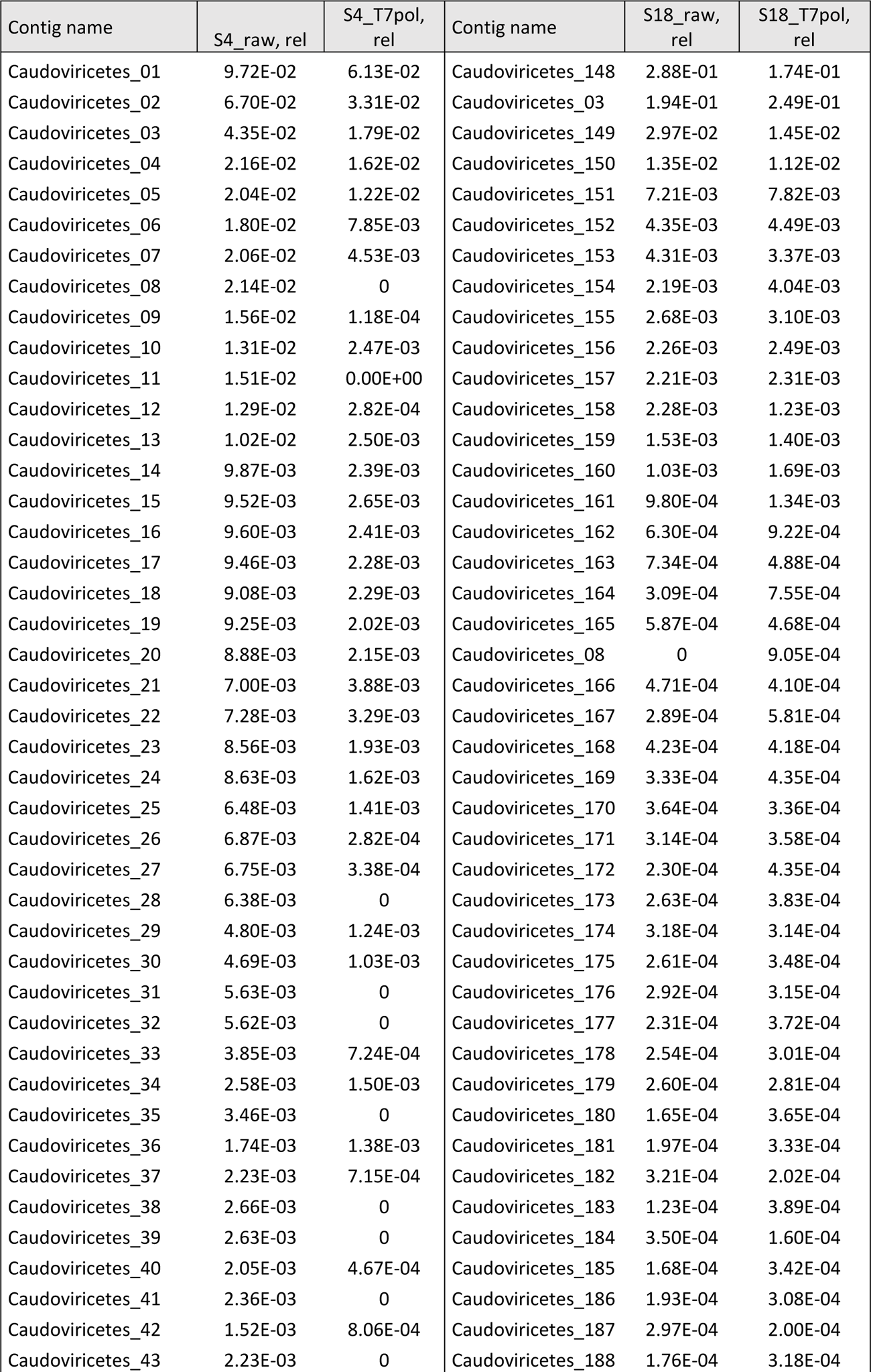
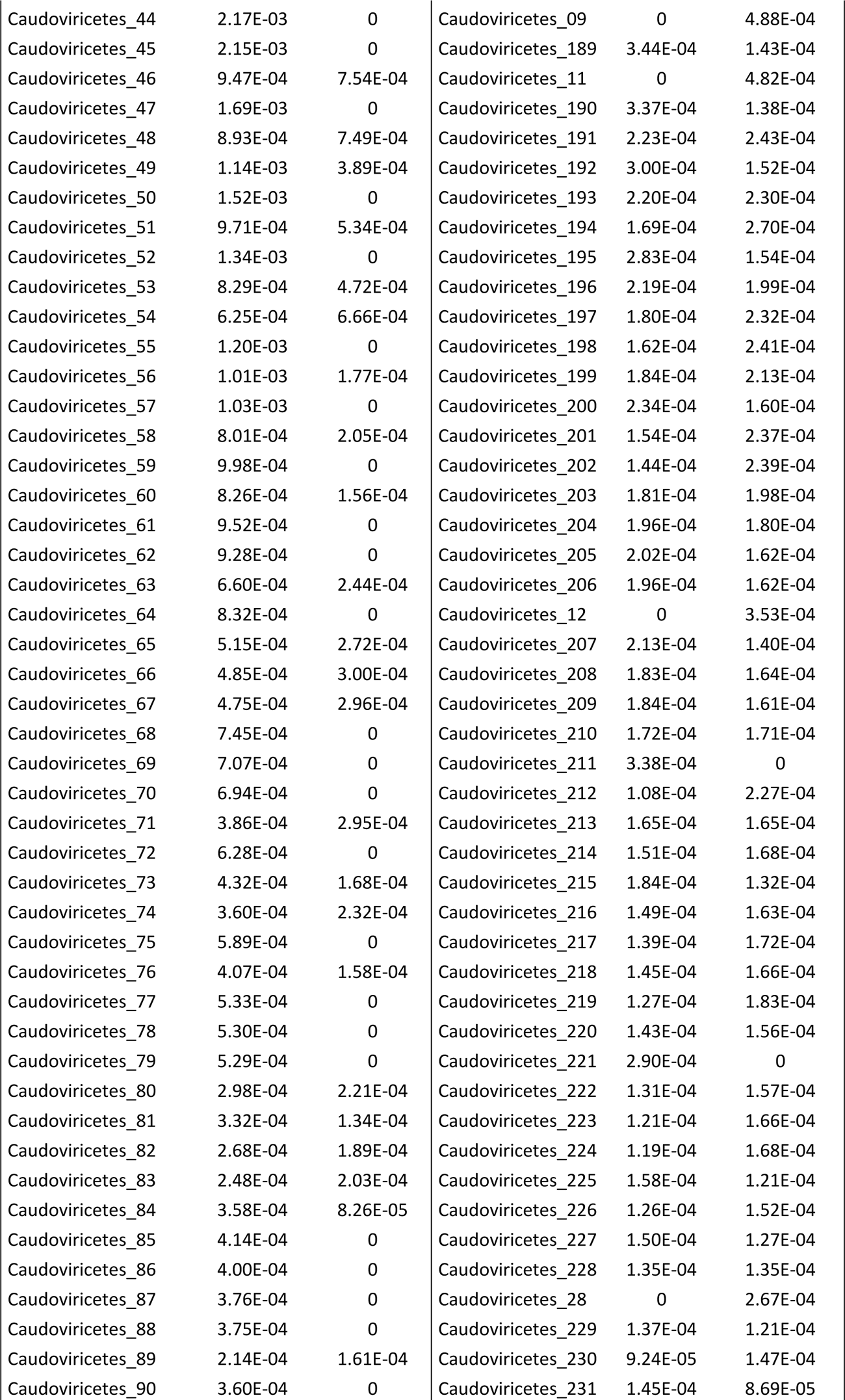
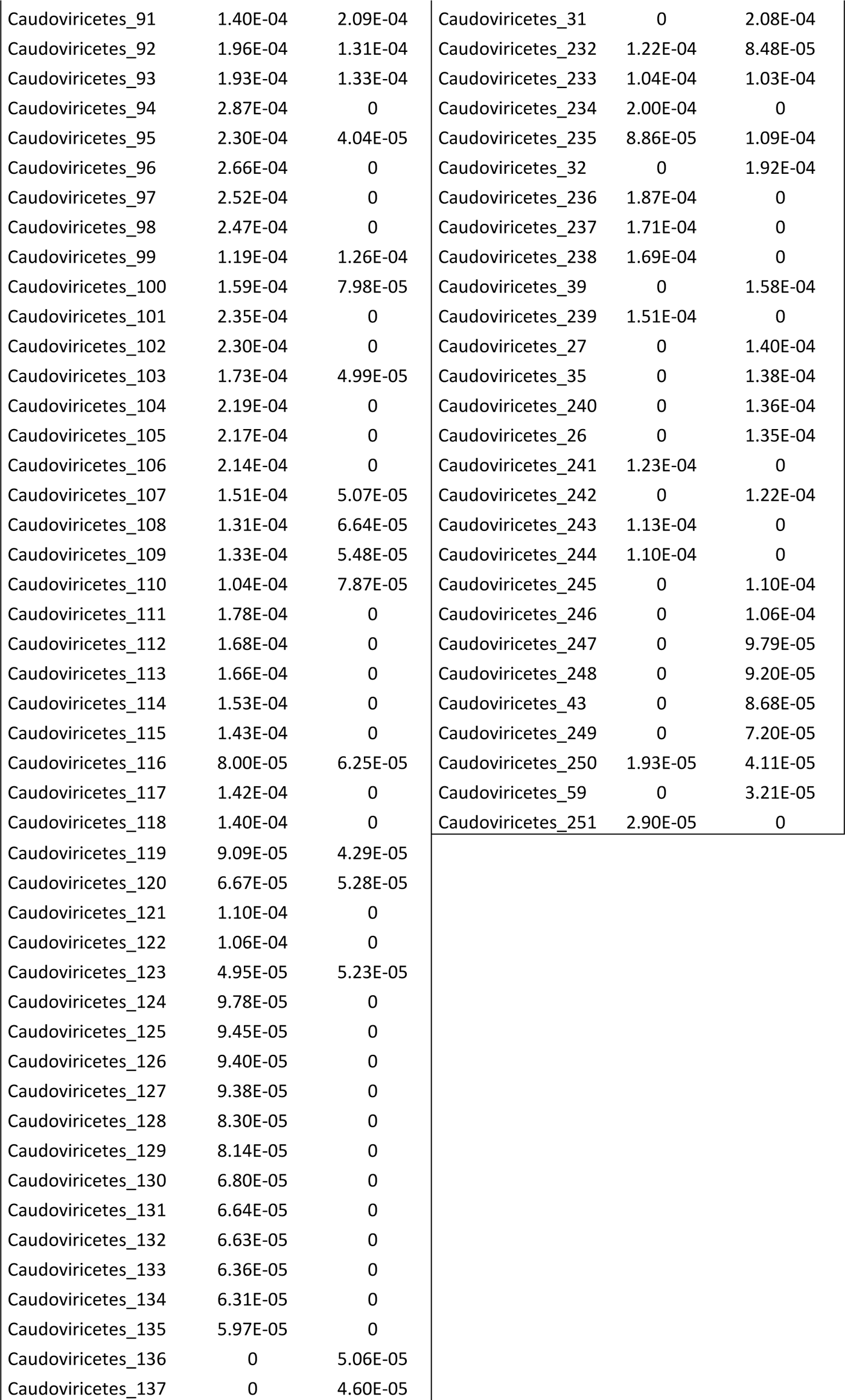
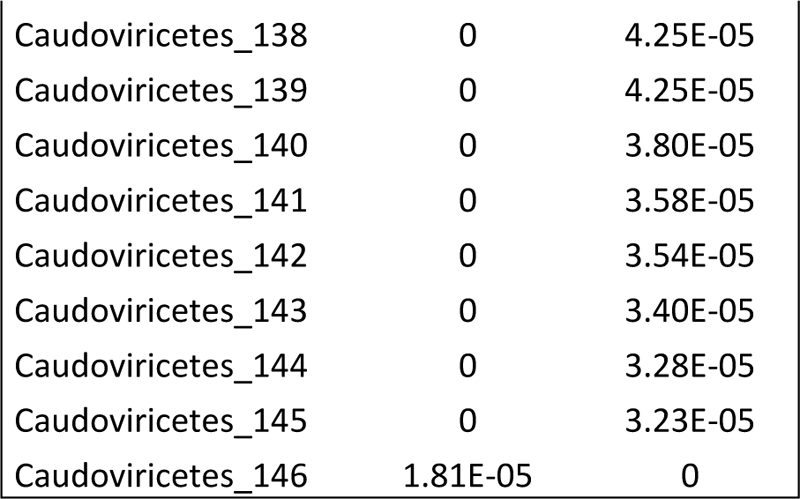
Relative abundance (rel) of each microviral contig in samples S4 and S18 before (“raw” column) and after T7pol treatment (“T7pol column”).

| Contig name | S4_raw, rel | S4_T7pol, rel | Contig name | S18_raw, rel | S18_T7pol, rel |
| --- | --- | --- | --- | --- | --- |
| Caudoviricetes_01 | 9.72E-02 | 6.13E-02 | Caudoviricetes_148 | 2.88E-01 | 1.74E-01 |
| Caudoviricetes_02 | 6.70E-02 | 3.31E-02 | Caudoviricetes_03 | 1.94E-01 | 2.49E-01 |
| Caudoviricetes_03 | 4.35E-02 | 1.79E-02 | Caudoviricetes_149 | 2.97E-02 | 1.45E-02 |
| Caudoviricetes_04 | 2.16E-02 | 1.62E-02 | Caudoviricetes_150 | 1.35E-02 | 1.12E-02 |
| Caudoviricetes_05 | 2.04E-02 | 1.22E-02 | Caudoviricetes_151 | 7.21E-03 | 7.82E-03 |
| Caudoviricetes_06 | 1.80E-02 | 7.85E-03 | Caudoviricetes_152 | 4.35E-03 | 4.49E-03 |
| Caudoviricetes_07 | 2.06E-02 | 4.53E-03 | Caudoviricetes_153 | 4.31E-03 | 3.37E-03 |
| Caudoviricetes_08 | 2.14E-02 | 0 | Caudoviricetes_154 | 2.19E-03 | 4.04E-03 |
| Caudoviricetes_09 | 1.56E-02 | 1.18E-04 | Caudoviricetes_155 | 2.68E-03 | 3.10E-03 |
| Caudoviricetes_10 | 1.31E-02 | 2.47E-03 | Caudoviricetes_156 | 2.26E-03 | 2.49E-03 |
| Caudoviricetes_11 | 1.51E-02 | 0.00E+00 | Caudoviricetes_157 | 2.21E-03 | 2.31E-03 |
| Caudoviricetes_12 | 1.29E-02 | 2.82E-04 | Caudoviricetes_158 | 2.28E-03 | 1.23E-03 |
| Caudoviricetes_13 | 1.02E-02 | 2.50E-03 | Caudoviricetes_159 | 1.53E-03 | 1.40E-03 |
| Caudoviricetes_14 | 9.87E-03 | 2.39E-03 | Caudoviricetes_160 | 1.03E-03 | 1.69E-03 |
| Caudoviricetes_15 | 9.52E-03 | 2.65E-03 | Caudoviricetes_161 | 9.80E-04 | 1.34E-03 |
| Caudoviricetes_16 | 9.60E-03 | 2.41E-03 | Caudoviricetes_162 | 6.30E-04 | 9.22E-04 |
| Caudoviricetes_17 | 9.46E-03 | 2.28E-03 | Caudoviricetes_163 | 7.34E-04 | 4.88E-04 |
| Caudoviricetes_18 | 9.08E-03 | 2.29E-03 | Caudoviricetes_164 | 3.09E-04 | 7.55E-04 |
| Caudoviricetes_19 | 9.25E-03 | 2.02E-03 | Caudoviricetes_165 | 5.87E-04 | 4.68E-04 |
| Caudoviricetes_20 | 8.88E-03 | 2.15E-03 | Caudoviricetes_08 | 0 | 9.05E-04 |
| Caudoviricetes_21 | 7.00E-03 | 3.88E-03 | Caudoviricetes_166 | 4.71E-04 | 4.10E-04 |
| Caudoviricetes_22 | 7.28E-03 | 3.29E-03 | Caudoviricetes_167 | 2.89E-04 | 5.81E-04 |
| Caudoviricetes_23 | 8.56E-03 | 1.93E-03 | Caudoviricetes_168 | 4.23E-04 | 4.18E-04 |
| Caudoviricetes_24 | 8.63E-03 | 1.62E-03 | Caudoviricetes_169 | 3.33E-04 | 4.35E-04 |
| Caudoviricetes_25 | 6.48E-03 | 1.41E-03 | Caudoviricetes_170 | 3.64E-04 | 3.36E-04 |
| Caudoviricetes_26 | 6.87E-03 | 2.82E-04 | Caudoviricetes_171 | 3.14E-04 | 3.58E-04 |
| Caudoviricetes_27 | 6.75E-03 | 3.38E-04 | Caudoviricetes_172 | 2.30E-04 | 4.35E-04 |
| Caudoviricetes_28 | 6.38E-03 | 0 | Caudoviricetes_173 | 2.63E-04 | 3.83E-04 |
| Caudoviricetes_29 | 4.80E-03 | 1.24E-03 | Caudoviricetes_174 | 3.18E-04 | 3.14E-04 |
| Caudoviricetes_30 | 4.69E-03 | 1.03E-03 | Caudoviricetes_175 | 2.61E-04 | 3.48E-04 |
| Caudoviricetes_31 | 5.63E-03 | 0 | Caudoviricetes_176 | 2.92E-04 | 3.15E-04 |
| Caudoviricetes_32 | 5.62E-03 | 0 | Caudoviricetes_177 | 2.31E-04 | 3.72E-04 |
| Caudoviricetes_33 | 3.85E-03 | 7.24E-04 | Caudoviricetes_178 | 2.54E-04 | 3.01E-04 |
| Caudoviricetes_34 | 2.58E-03 | 1.50E-03 | Caudoviricetes_179 | 2.60E-04 | 2.81E-04 |
| Caudoviricetes_35 | 3.46E-03 | 0 | Caudoviricetes_180 | 1.65E-04 | 3.65E-04 |
| Caudoviricetes_36 | 1.74E-03 | 1.38E-03 | Caudoviricetes_181 | 1.97E-04 | 3.33E-04 |
| Caudoviricetes_37 | 2.23E-03 | 7.15E-04 | Caudoviricetes_182 | 3.21E-04 | 2.02E-04 |
| Caudoviricetes_38 | 2.66E-03 | 0 | Caudoviricetes_183 | 1.23E-04 | 3.89E-04 |
| Caudoviricetes_39 | 2.63E-03 | 0 | Caudoviricetes_184 | 3.50E-04 | 1.60E-04 |
| Caudoviricetes_40 | 2.05E-03 | 4.67E-04 | Caudoviricetes_185 | 1.68E-04 | 3.42E-04 |
| Caudoviricetes_41 | 2.36E-03 | 0 | Caudoviricetes_186 | 1.93E-04 | 3.08E-04 |
| Caudoviricetes_42 | 1.52E-03 | 8.06E-04 | Caudoviricetes_187 | 2.97E-04 | 2.00E-04 |
| Caudoviricetes_43 | 2.23E-03 | 0 | Caudoviricetes_188 | 1.76E-04 | 3.18E-04 |
| Caudoviricetes_44 | 2.17E-03 | 0 | Caudoviricetes_09 | 0 | 4.88E-04 |
| Caudoviricetes_45 | 2.15E-03 | 0 | Caudoviricetes_189 | 3.44E-04 | 1.43E-04 |
| Caudoviricetes_46 | 9.47E-04 | 7.54E-04 | Caudoviricetes_11 | 0 | 4.82E-04 |
| Caudoviricetes_47 | 1.69E-03 | 0 | Caudoviricetes_190 | 3.37E-04 | 1.38E-04 |
| Caudoviricetes_48 | 8.93E-04 | 7.49E-04 | Caudoviricetes_191 | 2.23E-04 | 2.43E-04 |
| Caudoviricetes_49 | 1.14E-03 | 3.89E-04 | Caudoviricetes_192 | 3.00E-04 | 1.52E-04 |
| Caudoviricetes_50 | 1.52E-03 | 0 | Caudoviricetes_193 | 2.20E-04 | 2.30E-04 |
| Caudoviricetes_51 | 9.71E-04 | 5.34E-04 | Caudoviricetes_194 | 1.69E-04 | 2.70E-04 |
| Caudoviricetes_52 | 1.34E-03 | 0 | Caudoviricetes_195 | 2.83E-04 | 1.54E-04 |
| Caudoviricetes_53 | 8.29E-04 | 4.72E-04 | Caudoviricetes_196 | 2.19E-04 | 1.99E-04 |
| Caudoviricetes_54 | 6.25E-04 | 6.66E-04 | Caudoviricetes_197 | 1.80E-04 | 2.32E-04 |
| Caudoviricetes_55 | 1.20E-03 | 0 | Caudoviricetes_198 | 1.62E-04 | 2.41E-04 |
| Caudoviricetes_56 | 1.01E-03 | 1.77E-04 | Caudoviricetes_199 | 1.84E-04 | 2.13E-04 |
| Caudoviricetes_57 | 1.03E-03 | 0 | Caudoviricetes_200 | 2.34E-04 | 1.60E-04 |
| Caudoviricetes_58 | 8.01E-04 | 2.05E-04 | Caudoviricetes_201 | 1.54E-04 | 2.37E-04 |
| Caudoviricetes_59 | 9.98E-04 | 0 | Caudoviricetes_202 | 1.44E-04 | 2.39E-04 |
| Caudoviricetes_60 | 8.26E-04 | 1.56E-04 | Caudoviricetes_203 | 1.81E-04 | 1.98E-04 |
| Caudoviricetes_61 | 9.52E-04 | 0 | Caudoviricetes_204 | 1.96E-04 | 1.80E-04 |
| Caudoviricetes_62 | 9.28E-04 | 0 | Caudoviricetes_205 | 2.02E-04 | 1.62E-04 |
| Caudoviricetes_63 | 6.60E-04 | 2.44E-04 | Caudoviricetes_206 | 1.96E-04 | 1.62E-04 |
| Caudoviricetes_64 | 8.32E-04 | 0 | Caudoviricetes_12 | 0 | 3.53E-04 |
| Caudoviricetes_65 | 5.15E-04 | 2.72E-04 | Caudoviricetes_207 | 2.13E-04 | 1.40E-04 |
| Caudoviricetes_66 | 4.85E-04 | 3.00E-04 | Caudoviricetes_208 | 1.83E-04 | 1.64E-04 |
| Caudoviricetes_67 | 4.75E-04 | 2.96E-04 | Caudoviricetes_209 | 1.84E-04 | 1.61E-04 |
| Caudoviricetes_68 | 7.45E-04 | 0 | Caudoviricetes_210 | 1.72E-04 | 1.71E-04 |
| Caudoviricetes_69 | 7.07E-04 | 0 | Caudoviricetes_211 | 3.38E-04 | 0 |
| Caudoviricetes_70 | 6.94E-04 | 0 | Caudoviricetes_212 | 1.08E-04 | 2.27E-04 |
| Caudoviricetes_71 | 3.86E-04 | 2.95E-04 | Caudoviricetes_213 | 1.65E-04 | 1.65E-04 |
| Caudoviricetes_72 | 6.28E-04 | 0 | Caudoviricetes_214 | 1.51E-04 | 1.68E-04 |
| Caudoviricetes_73 | 4.32E-04 | 1.68E-04 | Caudoviricetes_215 | 1.84E-04 | 1.32E-04 |
| Caudoviricetes_74 | 3.60E-04 | 2.32E-04 | Caudoviricetes_216 | 1.49E-04 | 1.63E-04 |
| Caudoviricetes_75 | 5.89E-04 | 0 | Caudoviricetes_217 | 1.39E-04 | 1.72E-04 |
| Caudoviricetes_76 | 4.07E-04 | 1.58E-04 | Caudoviricetes_218 | 1.45E-04 | 1.66E-04 |
| Caudoviricetes_77 | 5.33E-04 | 0 | Caudoviricetes_219 | 1.27E-04 | 1.83E-04 |
| Caudoviricetes_78 | 5.30E-04 | 0 | Caudoviricetes_220 | 1.43E-04 | 1.56E-04 |
| Caudoviricetes_79 | 5.29E-04 | 0 | Caudoviricetes_221 | 2.90E-04 | 0 |
| Caudoviricetes_80 | 2.98E-04 | 2.21E-04 | Caudoviricetes_222 | 1.31E-04 | 1.57E-04 |
| Caudoviricetes_81 | 3.32E-04 | 1.34E-04 | Caudoviricetes_223 | 1.21E-04 | 1.66E-04 |
| Caudoviricetes_82 | 2.68E-04 | 1.89E-04 | Caudoviricetes_224 | 1.19E-04 | 1.68E-04 |
| Caudoviricetes_83 | 2.48E-04 | 2.03E-04 | Caudoviricetes_225 | 1.58E-04 | 1.21E-04 |
| Caudoviricetes_84 | 3.58E-04 | 8.26E-05 | Caudoviricetes_226 | 1.26E-04 | 1.52E-04 |
| Caudoviricetes_85 | 4.14E-04 | 0 | Caudoviricetes_227 | 1.50E-04 | 1.27E-04 |
| Caudoviricetes_86 | 4.00E-04 | 0 | Caudoviricetes_228 | 1.35E-04 | 1.35E-04 |
| Caudoviricetes_87 | 3.76E-04 | 0 | Caudoviricetes_28 | 0 | 2.67E-04 |
| Caudoviricetes_88 | 3.75E-04 | 0 | Caudoviricetes_229 | 1.37E-04 | 1.21E-04 |
| Caudoviricetes_89 | 2.14E-04 | 1.61E-04 | Caudoviricetes_230 | 9.24E-05 | 1.47E-04 |
| Caudoviricetes_90 | 3.60E-04 | 0 | Caudoviricetes_231 | 1.45E-04 | 8.69E-05 |
| Caudoviricetes_91 | 1.40E-04 | 2.09E-04 | Caudoviricetes_31 | 0 | 2.08E-04 |
| Caudoviricetes_92 | 1.96E-04 | 1.31E-04 | Caudoviricetes_232 | 1.22E-04 | 8.48E-05 |
| Caudoviricetes_93 | 1.93E-04 | 1.33E-04 | Caudoviricetes_233 | 1.04E-04 | 1.03E-04 |
| Caudoviricetes_94 | 2.87E-04 | 0 | Caudoviricetes_234 | 2.00E-04 | 0 |
| Caudoviricetes_95 | 2.30E-04 | 4.04E-05 | Caudoviricetes_235 | 8.86E-05 | 1.09E-04 |
| Caudoviricetes_96 | 2.66E-04 | 0 | Caudoviricetes_32 | 0 | 1.92E-04 |
| Caudoviricetes_97 | 2.52E-04 | 0 | Caudoviricetes_236 | 1.87E-04 | 0 |
| Caudoviricetes_98 | 2.47E-04 | 0 | Caudoviricetes_237 | 1.71E-04 | 0 |
| Caudoviricetes_99 | 1.19E-04 | 1.26E-04 | Caudoviricetes_238 | 1.69E-04 | 0 |
| Caudoviricetes_100 | 1.59E-04 | 7.98E-05 | Caudoviricetes_39 | 0 | 1.58E-04 |
| Caudoviricetes_101 | 2.35E-04 | 0 | Caudoviricetes_239 | 1.51E-04 | 0 |
| Caudoviricetes_102 | 2.30E-04 | 0 | Caudoviricetes_27 | 0 | 1.40E-04 |
| Caudoviricetes_103 | 1.73E-04 | 4.99E-05 | Caudoviricetes_35 | 0 | 1.38E-04 |
| Caudoviricetes_104 | 2.19E-04 | 0 | Caudoviricetes_240 | 0 | 1.36E-04 |
| Caudoviricetes_105 | 2.17E-04 | 0 | Caudoviricetes_26 | 0 | 1.35E-04 |
| Caudoviricetes_106 | 2.14E-04 | 0 | Caudoviricetes_241 | 1.23E-04 | 0 |
| Caudoviricetes_107 | 1.51E-04 | 5.07E-05 | Caudoviricetes_242 | 0 | 1.22E-04 |
| Caudoviricetes_108 | 1.31E-04 | 6.64E-05 | Caudoviricetes_243 | 1.13E-04 | 0 |
| Caudoviricetes_109 | 1.33E-04 | 5.48E-05 | Caudoviricetes_244 | 1.10E-04 | 0 |
| Caudoviricetes_110 | 1.04E-04 | 7.87E-05 | Caudoviricetes_245 | 0 | 1.10E-04 |
| Caudoviricetes_111 | 1.78E-04 | 0 | Caudoviricetes_246 | 0 | 1.06E-04 |
| Caudoviricetes_112 | 1.68E-04 | 0 | Caudoviricetes_247 | 0 | 9.79E-05 |
| Caudoviricetes_113 | 1.66E-04 | 0 | Caudoviricetes_248 | 0 | 9.20E-05 |
| Caudoviricetes_114 | 1.53E-04 | 0 | Caudoviricetes_43 | 0 | 8.68E-05 |
| Caudoviricetes_115 | 1.43E-04 | 0 | Caudoviricetes_249 | 0 | 7.20E-05 |
| Caudoviricetes_116 | 8.00E-05 | 6.25E-05 | Caudoviricetes_250 | 1.93E-05 | 4.11E-05 |
| Caudoviricetes_117 | 1.42E-04 | 0 | Caudoviricetes_59 | 0 | 3.21E-05 |
| Caudoviricetes_118 | 1.40E-04 | 0 | Caudoviricetes_251 | 2.90E-05 | 0 |
| Caudoviricetes_119 | 9.09E-05 | 4.29E-05 |  |  |  |
| Caudoviricetes_120 | 6.67E-05 | 5.28E-05 |  |  |  |
| Caudoviricetes_121 | 1.10E-04 | 0 |  |  |  |
| Caudoviricetes_122 | 1.06E-04 | 0 |  |  |  |
| Caudoviricetes_123 | 4.95E-05 | 5.23E-05 |  |  |  |
| Caudoviricetes_124 | 9.78E-05 | 0 |  |  |  |
| Caudoviricetes_125 | 9.45E-05 | 0 |  |  |  |
| Caudoviricetes_126 | 9.40E-05 | 0 |  |  |  |
| Caudoviricetes_127 | 9.38E-05 | 0 |  |  |  |
| Caudoviricetes_128 | 8.30E-05 | 0 |  |  |  |
| Caudoviricetes_129 | 8.14E-05 | 0 |  |  |  |
| Caudoviricetes_130 | 6.80E-05 | 0 |  |  |  |
| Caudoviricetes_131 | 6.64E-05 | 0 |  |  |  |
| Caudoviricetes_132 | 6.63E-05 | 0 |  |  |  |
| Caudoviricetes_133 | 6.36E-05 | 0 |  |  |  |
| Caudoviricetes_134 | 6.31E-05 | 0 |  |  |  |
| Caudoviricetes_135 | 5.97E-05 | 0 |  |  |  |
| Caudoviricetes_136 | 0 | 5.06E-05 |  |  |  |
| Caudoviricetes_137 | 0 | 4.60E-05 |  |  |  |

|  |  |  |
| --- | --- | --- |
| Caudoviricetes_138 | 0 | 4.25E-05 |
| Caudoviricetes_139 | 0 | 4.25E-05 |
| Caudoviricetes_140 | 0 | 3.80E-05 |
| Caudoviricetes_141 | 0 | 3.58E-05 |
| Caudoviricetes_142 | 0 | 3.54E-05 |
| Caudoviricetes_143 | 0 | 3.40E-05 |
| Caudoviricetes_144 | 0 | 3.28E-05 |
| Caudoviricetes_145 | 0 | 3.23E-05 |
| Caudoviricetes_146 | 1.81E-05 | 0 |

**Supplementary figure 1:**
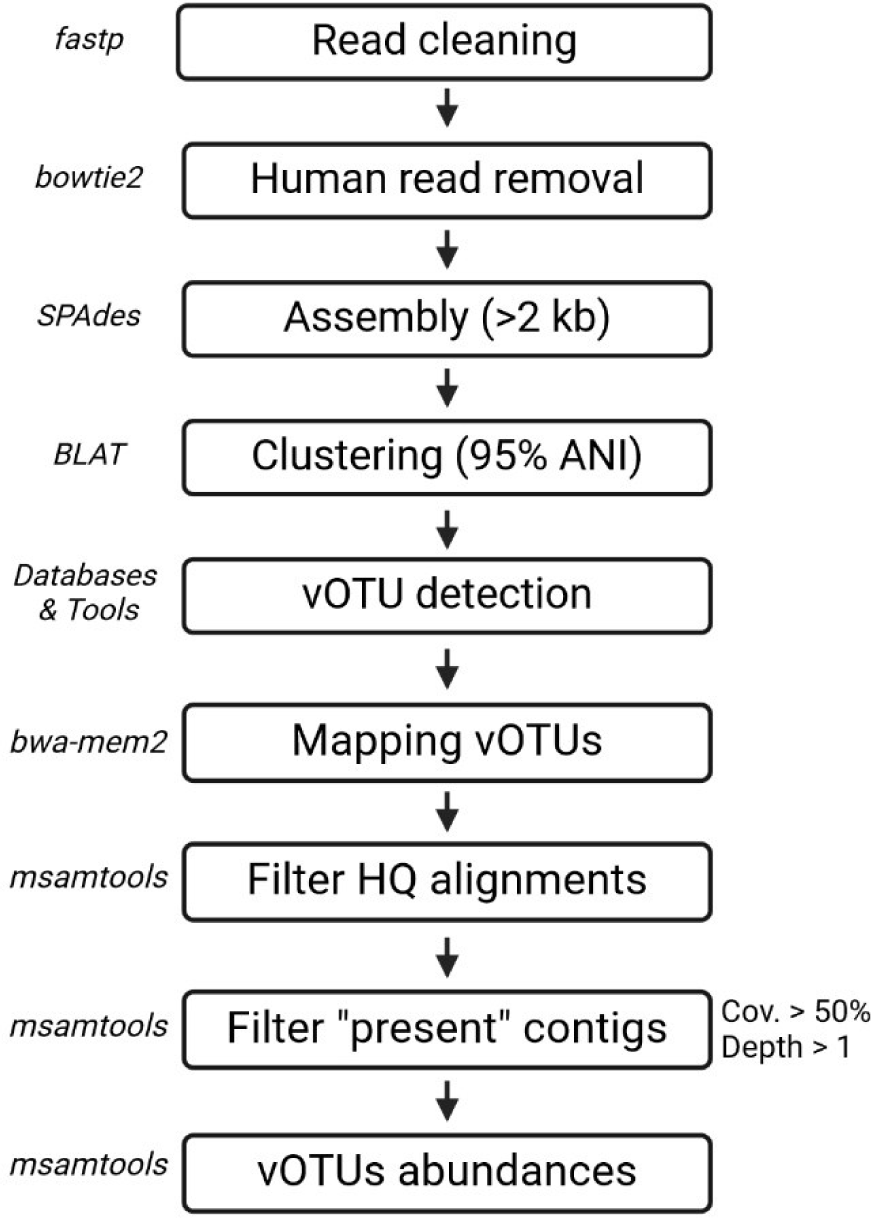
pipeline of virome reads analysis

**Supplementary figure 2:**
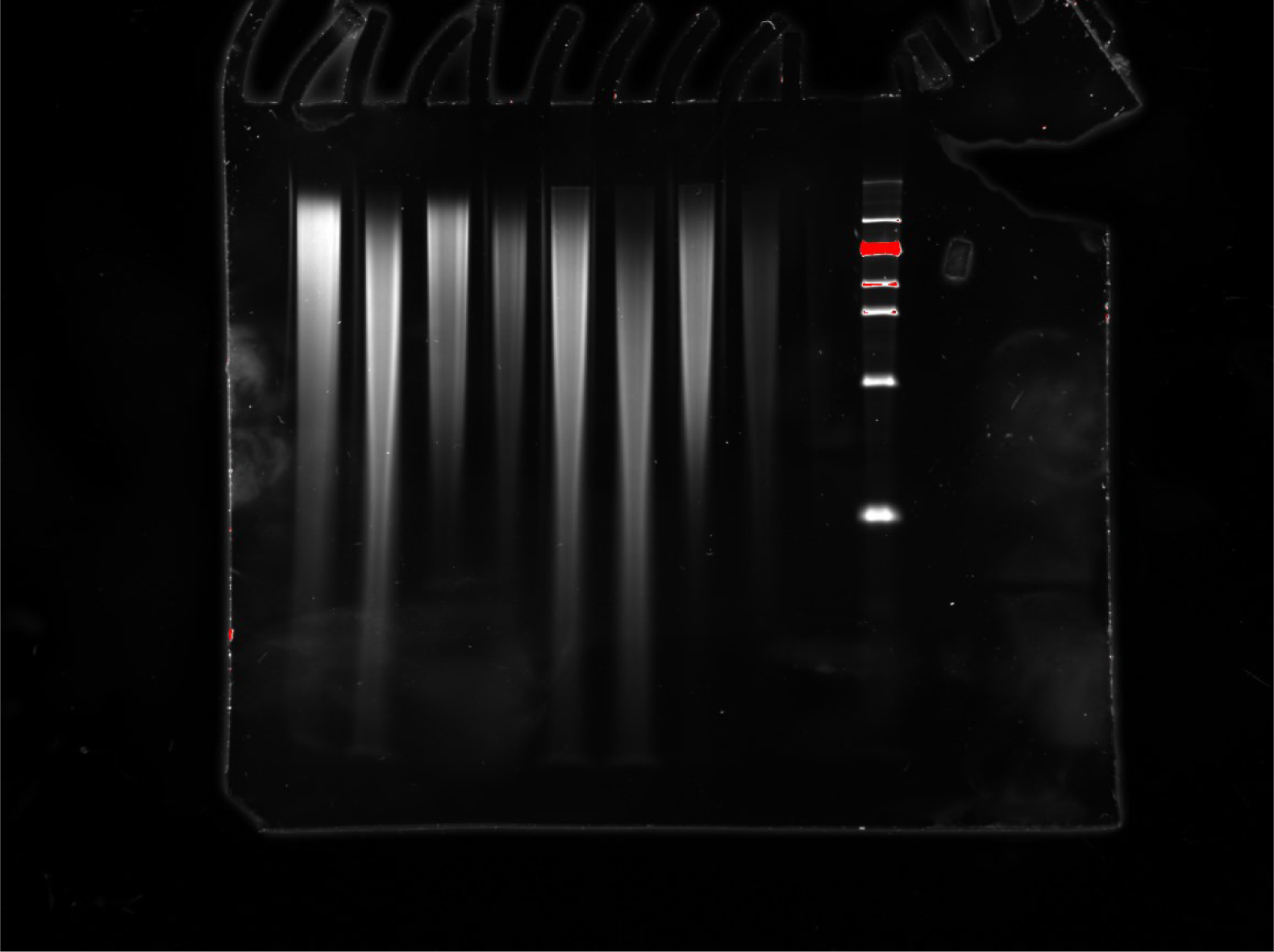
raw gel of figure 1

**Supplementary figure 3:**
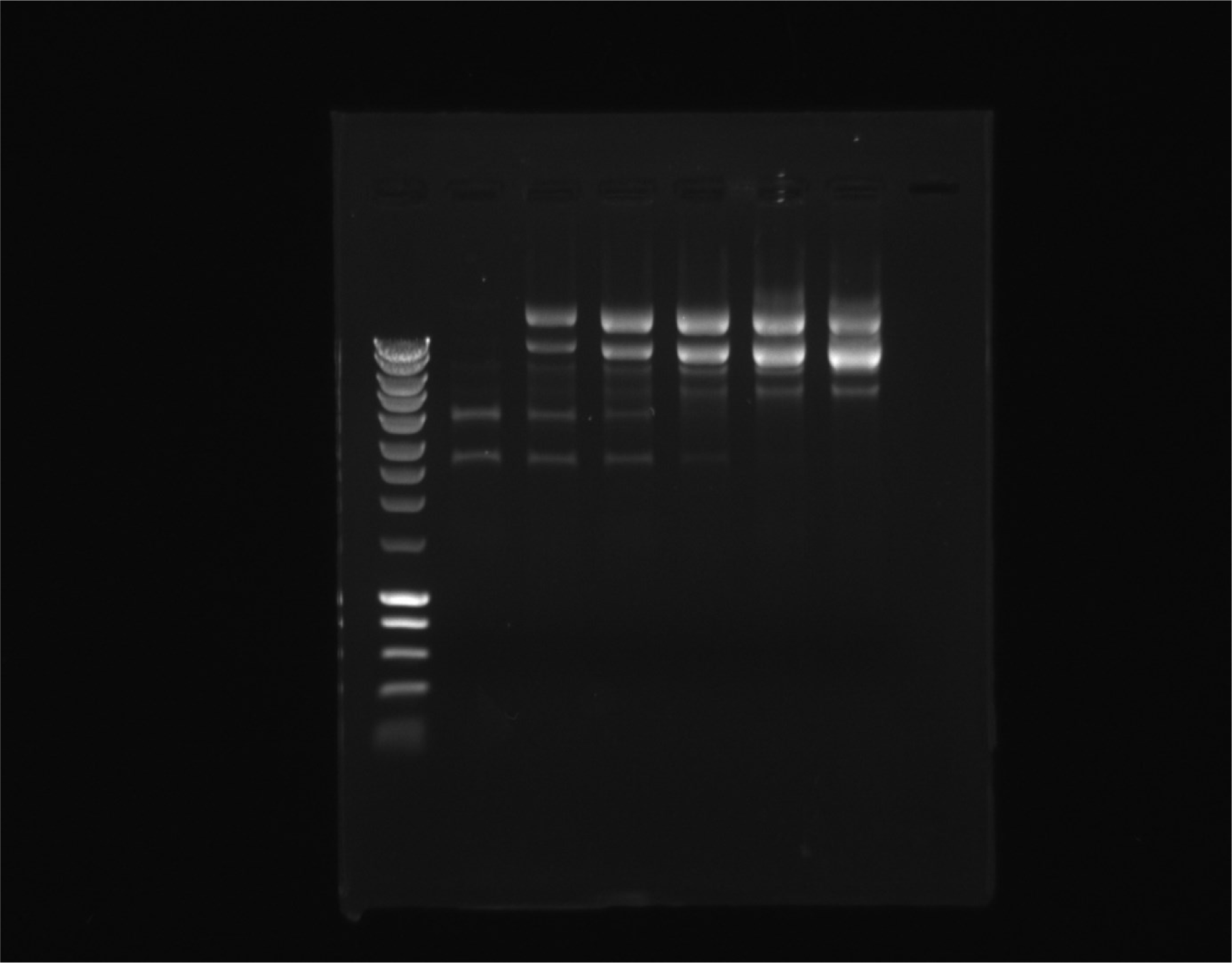
raw gel of figure 2.

**Supplementary figure 4:**
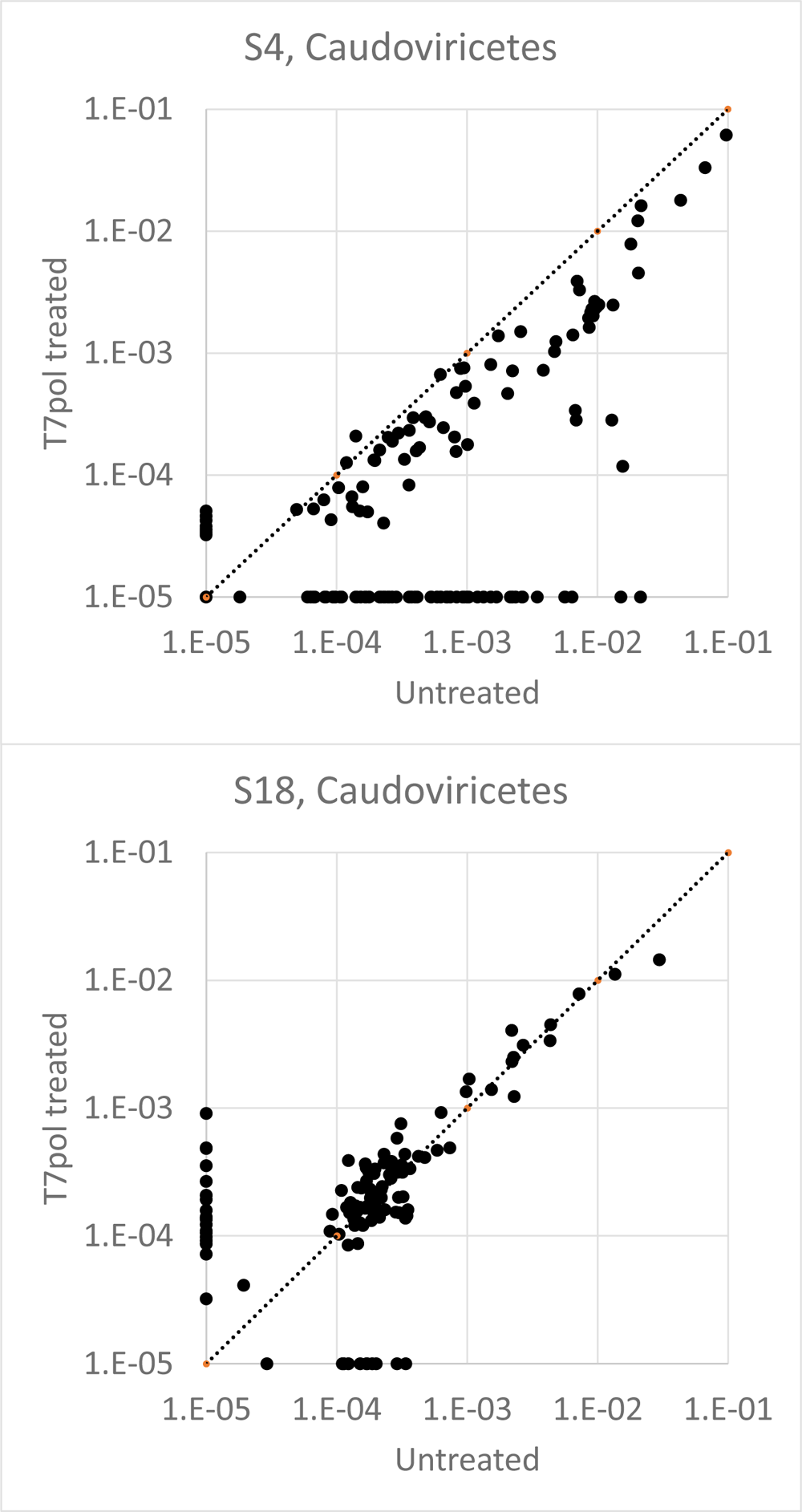
Relative abundance of each *Caudoviricetes* detected in samples S4 (top) and S18 (bottom). Each dot represents one contig, with its abundance relative to all phages of the sample when sequenced without prior treatment for ssDNA (X axis) or after T7pol treatment (Y axis). The dotted line shows the bisector. Contigs undetected with one of the treatments were assigned an artificial value of 10^-5^.

**Supplementary figure 5:**
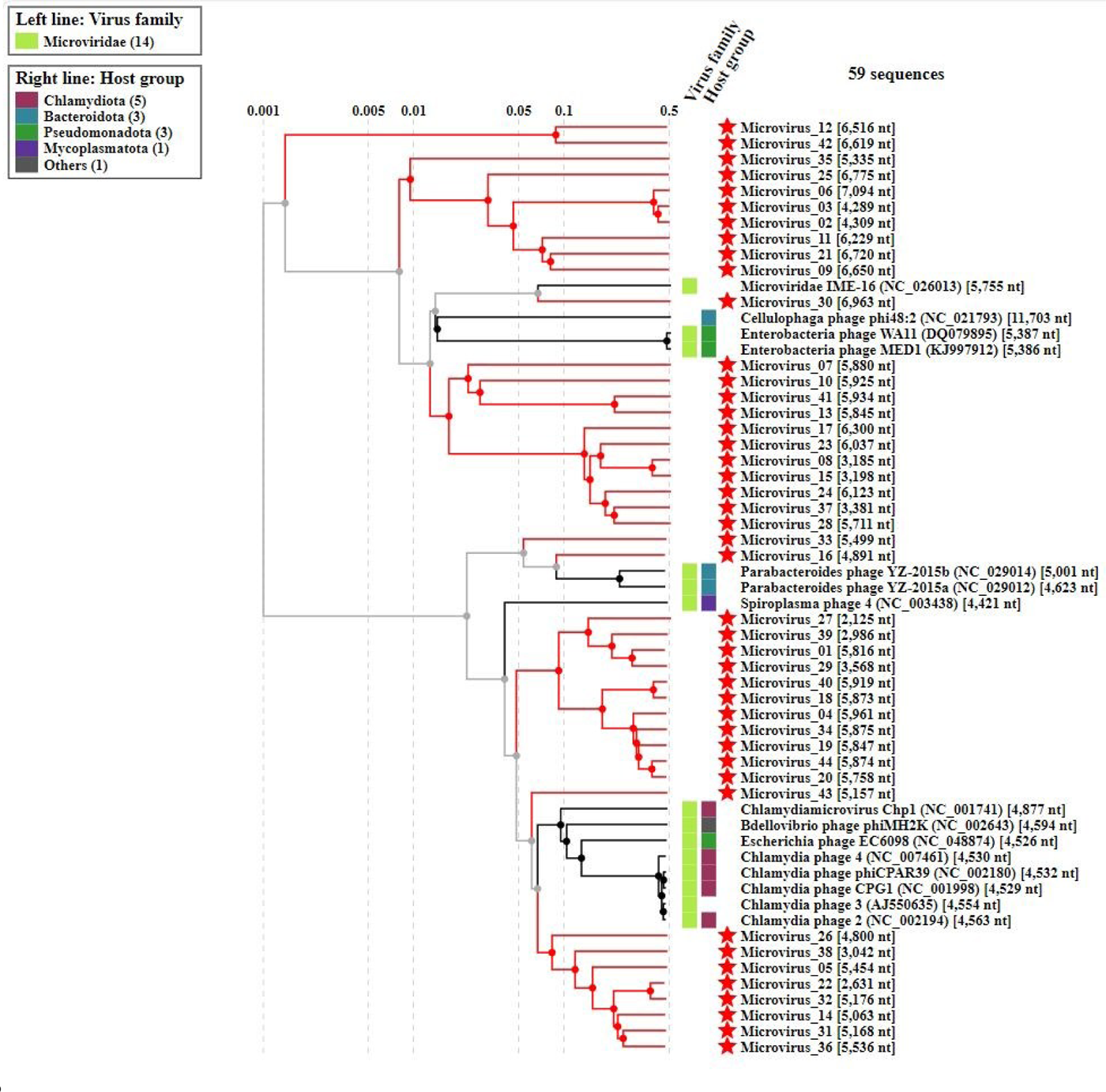
VipTree classification of the microviruses detected in samples S4 and S18.

